# Super-resolution live-cell mapping of protein-protein interactions using chemogenetic split reporters and STED microscopy

**DOI:** 10.1101/2025.08.05.668682

**Authors:** Stephanie Board, Arnaud Gautier

**Affiliations:** Sorbonne Université, École Normale Supérieure, Université PSL, CNRS, Chimie Physique et Chimie du Vivant (CPCV), 75005 Paris, France; Institut Universitaire de France, Paris France

**Keywords:** Protein-protein interactions, super-resolution imaging, chemogenetic split fluorescent reporters, Stimulated Emission Depletion (STED) nanoscopy

## Abstract

The ability to map protein-protein interactions (PPI) within living cells at high spatial resolution is essential for unravelling their roles in cellular biology. Although several super-resolution microscopy techniques are available, visualizing PPI below the diffraction limit of light across entire live cells remains a challenge. Here, we introduce an approach that combines the chemogenetic split fluorescent reporter splitFAST2 with STED microscopy for sub-diffraction imaging of PPI. As the fluorescence of splitFAST2 is activated only where the two proteins interact, this system allows for highly precise and unambiguous localization in live cells, enhanced further by the rapid super-resolution imaging capabilities of STED. The improved spatial resolution of our approach enabled us to precisely map the subcellular localization of inducible interactions or constitutive interactions. Beyond simply detecting PPIs below the diffraction limit in whole living cells, we also demonstrated that splitFAST2 can serve as the basis for fluorescent probes with very low background, enabling effective subdiffraction imaging of repetitive cellular structures like filamentous actin and microtubules.

## INTRODUCTION

Protein-protein interactions (PPI) play an essential role in regulating and orchestrating most cellular processes. The functions of these interactions are tightly related to their spatial distribution and temporal dynamics, making direct visualization of PPIs at high resolution critical for understanding biological processes. Achieving high resolution imaging of PPI has become possible through advancements in super-resolution fluorescence microscopy techniques. These methods provides nanoscale insights into molecular distribution, significantly enhancing our understanding of cell biology^[1]^.

A variety of super-resolution techniques are now available, including single molecule localization microscopy (SMLM)^[2–4]^, stimulated emission depletion (STED) microscopy^[5,6]^ or MINFLUX^[7]^ that combined structured illumination and single molecule localization. While these tools can theoretically offer nanoscale visualization of PPIs through two-color co-localization, challenges remain as accurate analysis can be limited by background noise from non-interacting molecules and the inherent challenge of discerning truly co-localized fluorophores due to limited spatial resolution, even in sub-diffraction microscopy. Alternatively, Förster Resonance Energy Transfer (FRET), which relies on the highly distance-dependent efficiency of energy transfer between donor and acceptor fluorophores, offers an efficient approach to study and visualize PPI. However, although super-resolved FRET images have been obtained by combining FRET with STED^[8]^, achieving super-resolution FRET imaging remains technically challenging.

One effective method to decouple proximity detection from optical resolution is through bimolecular fluorescence complementation. This technique uses two non-fluorescent complementary fragments of a fluorescent reporter that reassemble upon interaction of two proteins, making their proximity visible. Leveraging this strategy, split photocontrollable fluorescent proteins have been developed for sub-diffraction imaging of PPI using super-resolution methods such as photoactivated fluorescence localization microscopy (PALM)^[3]^ or stochastic optical fluctuation imaging (SOFI). For example, splitting of green-to-red photoconvertible fluorescent proteins have enabled approaches like BiFC-PALM^[9,10]^, while split version of reversibly photoswitching green fluorescent proteins have been used in reconstituted fluorescence-based SOFI (refSOFI)^[11]^. Another innovation includes the split version of HaloTag (so-called TagBiFC), which allows the use of organic dyes with superior brightness and photostability for long-term single molecule tracking of protein-protein interactions^[12]^. While BiFC-PALM and TagBiFC can achieve impressive spatial resolution, these methods rely on the inherently slow acquisition of single-molecule localization microscopy. As a result, these approaches are generally limited to fixed cells, making it challenging to study dynamic PPIs in live cells.

In this study, we present a novel approach for sub-diffraction imaging of PPIs in live cells, which integrates the chemogenetic fluorescent reporter splitFAST2^[13]^ with STED nanoscopy (**Fig. 1**). Unlike SMLM methods relying on stochastic blinking of fluorophores, STED offers rapid imaging capabilities, making it particularly suited for whole-cell PPI visualization in live cells. Our work marks, to our knowledge, the first application of a split fluorescent reporter in combination with STED to achieve subdiffraction PPI imaging in live-cell environments.

**Figure 1.**
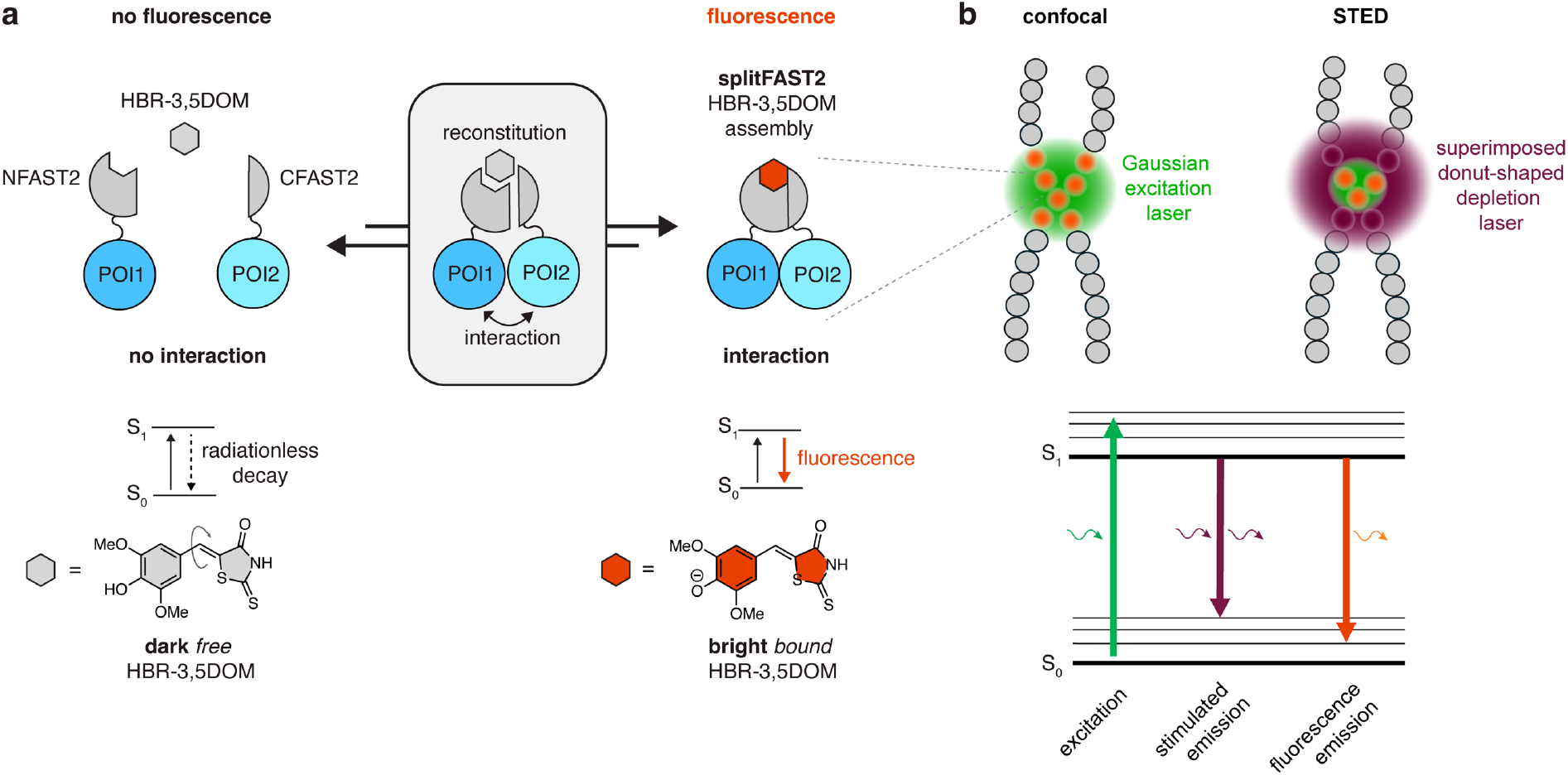
Subdiffraction imaging of protein-protein interaction in live cells using splitFAST2 and STED microscopy. **a** Principle of splitFAST2. Two complementary fragments, NFAST2 and CFAST2, reconstitute into a fluorescent assembly (in the presence of the exogenously applied fluorogenic chromophore HBR3,5DOM) upon interaction of two proteins (POI: protein of interest), making their proximity visible by fluorescence microscopy. **b** Principle of STED. The superimposition of a donut-shaped depletion laser at 775 nm to the Gaussian excitation laser enables to achieve sub-diffraction resolution. The depletion laser switches off fluorophores by stimulated emission in the donut region enabling to reduce the size of the point spread function, increasing thus the spatial resolution.

SplitFAST2 is an enhanced version of splitFAST, a chemogenetic fluorescent sensor designed to detect dynamic PPI with rapid and reversible complementation^[13,14]^. Unlike split fluorescent proteins and split-HaloTag, splitFAST and its derivatives stand out for their ability to visualize live-cell PPIs in real-time^[13–16]^ and track dynamic systems such as membrane contact sites^[17,18]^. SplitFAST was engineered by splitting the chemogenetic reporter FAST – a small protein of 14 kDa that binds and stabilizes the fluorescent state of fluorogenic 4-hydroxybenzylidene rhodanine chromophores that are otherwise non-fluorescent when unbound^[19]^. The enhanced splitFAST2 offers lower self-complementation, greater brightness, increased complementation efficiency and a higher dynamic range^[13]^. Previous studies showed that full-length FAST variants have favorable photophysical properties enabling efficient live-cell STED imaging^[20,21]^. One advantage of systems such as FAST and other fluorogen-activating proteins is their reversible, non-covalent fluorogen binding, which can participate to the photostability of the reporter^[22–24]^, enabling to sustain signal under high-intensity illumination through continuous exchange of bleached chromophores with new ones. Consequently, we expanded our investigation to splitFAST2 for sub-diffraction imaging of PPIs in live cells. We showed that this novel approach enables effective imaging of chemically induced and constitutively present PPIs with enhanced resolution in live cells, combining the advantages of PPI sensors with super-resolution imaging.

## RESULTS

To evaluate the performance of splitFAST2 and its suitability for STED, we first compared its photostability to that of full-length pFAST, previously shown to be compatible with STED imaging when paired with the fluorogenic chromophore HBR-3,5DOM^[20,21]^. The two complementary fragments of splitFAST2, CFAST2 (11 amino acids) and NFAST2 (114 amino acids), were fused at the C-terminus of FK506-binding protein (FKBP) and FKBP-rapamycin binding domain (FRB) of the mechanistic target of rapamycin (mTOR) respectively. We previously reported that induction of the FKBP-FRB interaction with the small molecule rapamycin enables efficient complementation of splitFAST2 in cells pre-treated with HBR-3,5DOM^[13]^. The FRB-NFAST2 fusion was further fused to the C-terminus of histone H2B for nuclear targeting to facilitate quantification. The two proteins H2B-FRB-NFAST2 and FKBP-CFAST2 were co-expressed in HeLa cells for 24 h before treatment with 10 μM HBR-3,5DOM and 100 nM rapamycin to preform splitFAST2. Long-term imaging by confocal microscopy exciting with a 560 nm laser allowed us to evaluate the photostability of splitFAST2 and compare it to that of pFAST imaged in the same conditions (**Supplementary Fig. 1**). Over a 600-frame time-lapse series, splitFAST2 retained approximately 60% of its initial fluorescence, while pFAST maintained around 85%. Although splitFAST2 displayed slightly lower photostability, it remained sufficiently photostable to support its potential compatibility with STED microscopy. Photostability comparison under STED imaging conditions yielded similar results, confirming the potential of splitFAST2 probes for STED imaging (**Supplementary Fig. 2)**.

To demonstrate the suitability of splitFAST2 for subdiffraction imaging of PPI, we designed sensors to detect chemically induced interactions at highly organized structures within cells, such as the microtubule network or organellar membranes from mitochondria. Specifically, we fused FRB-NFAST2 at the C-terminus of (i) the microtubule binding domain of the microtubule associated protein MAP4, and of (ii) the outer mitochondrial membrane targeting sequence of TOM20, a component of the translocase of the outer mitochondrial membrane TOM complex (**Supplementary Fig. 3**). When these constructs were co-expressed together with FKBP-CFAST2 in HeLa cells, rapamycin addition induced fluorescence complementation at the targeted subcellular structures, confirming the proximity-dependent complementation and high dynamic range of splitFAST2.

These constructs allowed us to induce splitFAST2 reconstitution at the microtubule network and mitochondrial membranes in live mammalian cells, and to record images using both conventional confocal microscopy and STED microscopy (**Fig. 2**). In the presence of HBR-3,5DOM, splitFAST2 emits fluorescence at approximatively 600 nm, offering a resolution limit of 300-400 nm laterally in conventional confocal microscopy. In STED, the superimposition of a donut-shaped depletion laser at 775 nm to the excitation laser enables to achieve sub-diffraction resolution (**Fig. 1**). The depletion laser switches off fluorophores by stimulated emission in the donut region enabling to reduce the size of the point spread function, increasing thus the spatial resolution. Using STED, we were able to increase twofold the global image resolution, enabling a clearer visualization of targeted subcellular structures, otherwise poorly resolved in diffraction-limited microscopy. The global image resolution enhancement over confocal microscopy was estimated using a parameter-free image resolution method based on image partial phase autocorrelation^[25]^. This resolution enhancement enabled us to successfully resolve microtubule filaments in tightly packed bundles (with full width at half-maximum as low as 70 nm) (**Fig. 2a-d**) and the membranes of mitochondria (**Fig. 2e-h**). The attained resolution closely matched that achieved when using full length pFAST reporter (**Supplementary Fig. 4**). This set of data demonstrated the effectiveness of splitFAST2 for sub-diffraction imaging of PPIs in live cells using STED microscopy.

**Figure 2.**
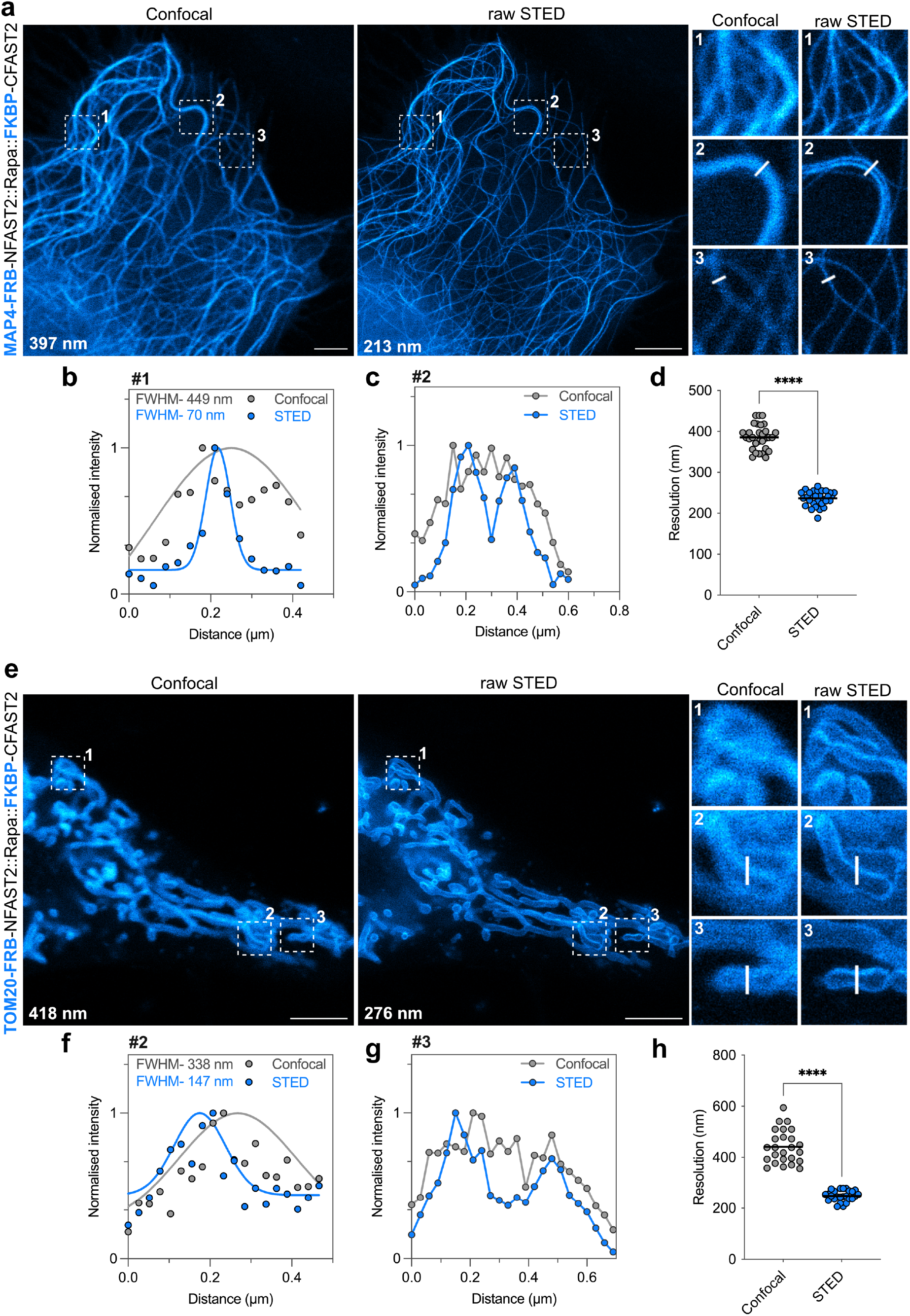
Live-cell subdiffraction STED imaging of inducible PPI. **a-d** Confocal and raw STED micrographs of live HeLa cells expressing MAP4-FRB-NFAST2 and FKBP-CFAST2 pretreated with 10 µM of HBR-3,5DOM. FRB-FKBP interaction was induced by addition of 100 nM rapamycin. Three regions of interest were selected for close-up comparison. Micrographs are representative of n = 30 cells from three independent experiments. Global image resolution is given on the image. Scale bars, 5 µm. **b**,**c** Line profile across microtubule filaments from close-up were used to compare gain in resolution between confocal and STED. Graph shows raw data of line profile from confocal and STED images (points), in addition to the corresponding gaussian fit (line) in the case of (**b**). **d** Comparison of resolution of confocal and STED micrographs using image decorrelation analysis^[25]^ (n = 30 cells). **e-h** Confocal and raw STED micrographs of live HeLa cells expressing TOM20-FRB-NFAST2 and FKBP-CFAST2 pretreated with 10 µM of HBR-3,5DOM. FRB-FKBP interaction was induced by addition of 100 nM rapamycin. Three regions of interest were selected for close-up comparison. Micrographs are representative of n = 25 cells from three independent experiments. Global image resolution is given on the image. Scale bars, 5 µm. **f**,**g** Line profile across mitochondria membranes from close-up were used to compare gain in resolution between confocal and STED. Graph shows raw data of line profile from confocal and STED images (points), in addition to the corresponding gaussian fit (line) in the case of (**f**). **h** Comparison of resolution of confocal and STED micrographs using image decorrelation analysis^[25]^ (n = 25 cells). See supplementary Table xxx for detailed acquisition parameters.

Building on these results, we further investigated the potential of splitFAST2 to visualize constitutive PPI. As a model interaction, we first focused on the direct interaction between filamentous actin (F-actin) and LifeAct, a small peptide of 17 amino acids from the yeast Abp140 protein that binds specifically to F-actin (**Supplementary Fig. 2** and **Fig. 3**). The genetic tagging of β-actin is often challenging due to the interference large tags can cause on actin polymerization^[26]^. Here, we benefited from the small size of CFAST2 to genetically tag β-actin with minimal perturbation. When NFAST2-LifeAct and CFAST2-β-actin were co-expressed in mammalian cells in presence of HBR-3,5DOM, splitFAST2 successfully complemented, confirming the interaction of LifeAct with F-actin (**Supplementary Fig. 5**). This set up revealed an intricate and dense intracellular actin network. The combination NFAST2-LifeAct and CFAST2-β-actin allowed the visualization of thin actin structures, including filopodia and specialized actin-containing formations like nuclear actin cages (**Supplementary Fig. 5**). By harnessing STED microscopy, we achieved sub-diffraction imaging of the intricate architecture of actin filaments in live cells (**Fig. 3** and **Supplementary Fig. 6**). This approach revealed detailed three-dimensional actin networks, including stress fibers and tightly bundled filaments within microvilli and filopodia. Notably, the finest structures – such as those in microvilli and filopodia – resolved with a full width at half-maximum of just 90 nm, a substantial improvement over conventional confocal microscopy.

**Figure 3.**
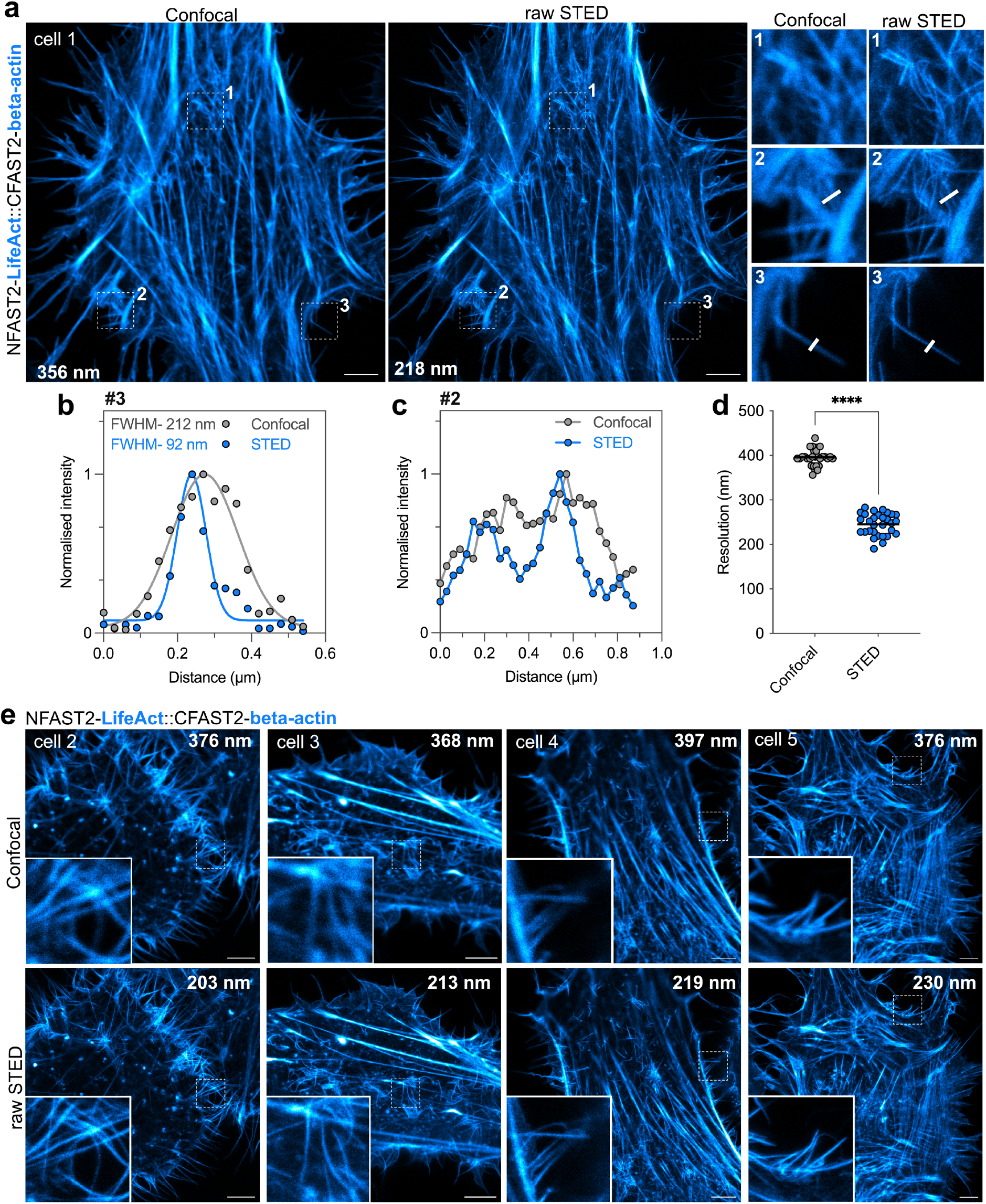
Live-cell subdiffraction STED imaging of LifeAct-F-actin interaction. **a**,**e** Confocal and raw STED micrographs of live HeLa cells expressing NFAST2-LifeAct and CFAST2-β-actin treated with 10 µM of HBR-3,5DOM. In (**a**) three regions of interest were selected for close-up comparison. Micrographs are representative of n = 30 cells from three independent experiments. Global image resolution is given on the image. Scale bars, 5 µm. **b**,**c** Line profile across actin filaments from close-up in (**a**) were used to compare gain in resolution between confocal and STED. Graph shows raw data of line profile from confocal and STED images (points), in addition to the corresponding gaussian fit (line) in the case of (**b**). **d** Comparison of resolution of confocal and STED micrographs using image decorrelation analysis^[25]^ (n= 30 cells). See supplementary Table xxx for detailed acquisition parameters.

Next, we coupled splitFAST2 and STED to examine the self-association of the type V intermediate filament protein Lamin A/C^[27]^ at sub-diffraction resolution (**Fig. 4a-c**). Lamin A/C assemble into the nuclear lamina through a hierarchical process^[27,28]^. Initially, parallel coiled-coil dimers are formed, which then associate head-to-tail into protofilaments. These protofilaments further engage in lateral association to create mature thick intermediate filaments. These filaments play a crucial role in maintaining the structural integrity of the nucleus, supporting the inner nuclear membrane, and contributing to nuclear shape, mechanical stability, and chromatin organization regulation. Fusion of the two halves of splitFAST2 to Lamin A/C enable the effective visualization of the Lamin A/C oligomerization in live cells (**Fig. 4a-c**). STED super-resolution imaging enabled a significant increase in effective resolution. When plotting intensity profiles of lines drawn perpendicular to the lamina in confocal and STED images, we found a clear difference between the two imaging techniques: the smallest FWHM measured by STED was 170 nm, while it was 323 nm for confocal microscopy.

**Figure 4.**
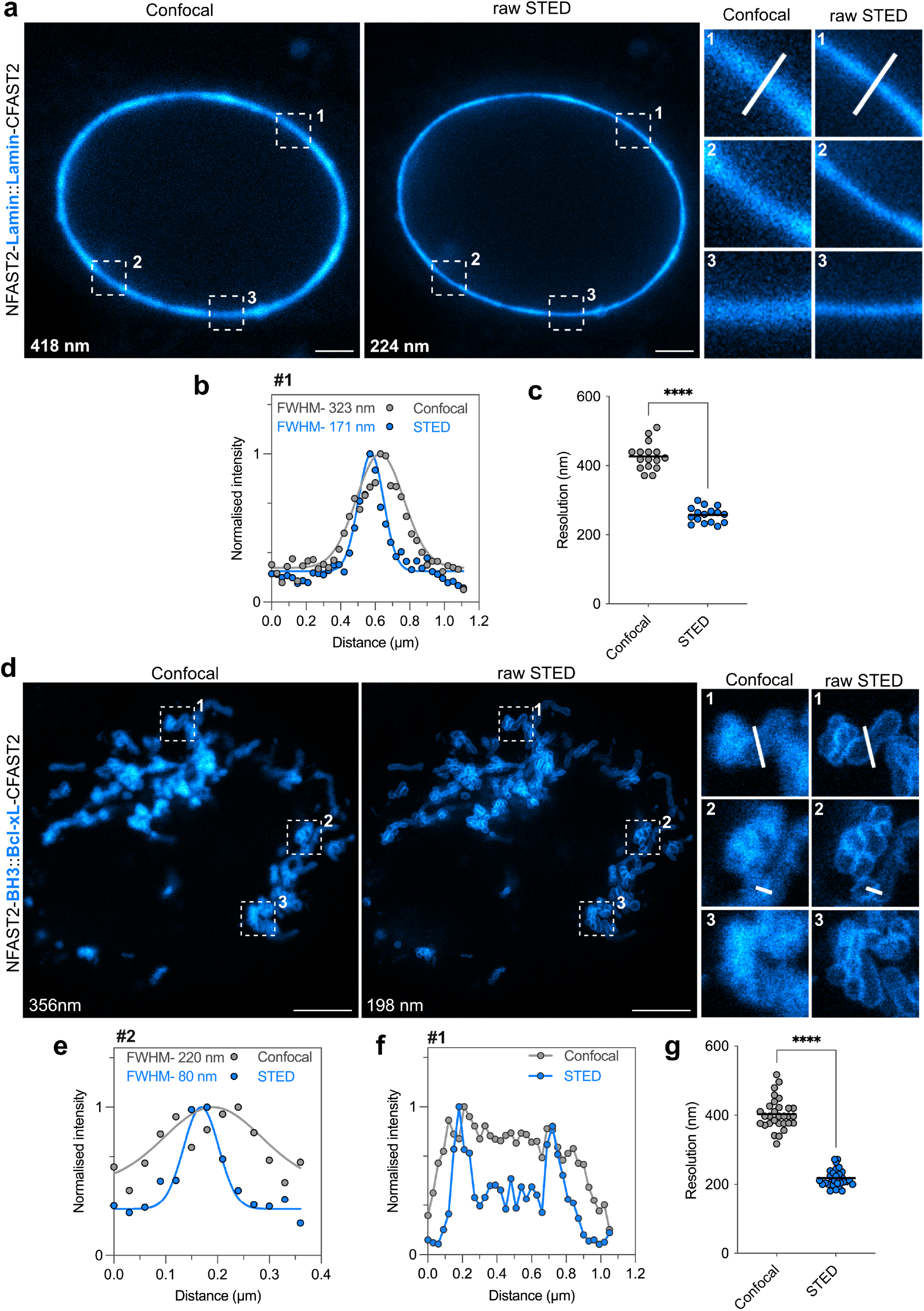
Live-cell subdiffraction STED imaging of constitutive PPI. **a-c** Confocal and raw STED micrographs of live HeLa cells expressing NFAST2-Lamin and CFAST2-Lamin treated with 10 µM of HBR-3,5DOM. Three regions of interest were selected for close-up comparison. Micrographs are representative of n = 15 cells from three independent experiments. Global image resolution is given on the image. Scale bars, 5 µm. **b** Line profile across nuclear membrane from close-up were used to compare gain in resolution between confocal and STED. Graph shows raw data of line profile from confocal and STED images (points), in addition to the corresponding gaussian fit (line). **c** Comparison of resolution of confocal and STED micrographs using image decorrelation analysis^[25]^ (n = 15 cells). **d-g** Confocal and raw STED micrographs of live HeLa cells expressing NFAST2-BH3 and Bcl-xL-CFAST2 treated with 10 µM of HBR-3,5DOM. Three regions of interest were selected for close-up comparison. Micrographs are representative of n = 30 cells from three independent experiments. Global image resolution is given on the image. Scale bars, 5 µm. e,f Line profile across mitochondria membranes from close-up were used to compare gain in resolution between confocal and STED. Graph shows raw data of line profile from confocal and STED images (points), in addition to the corresponding gaussian fit (line) in the case of (**e**). **g** Comparison of resolution of confocal and STED micrographs using image decorrelation analysis^[25]^ (n = 30 cells). See supplementary Table xxx for detailed acquisition parameters.

Finally, we extended the application of splitFAST2 for subdiffraction PPI imaging by utilizing STED microscopy to observe the interaction between Bcl-xL and its partners (**Fig. 4d-g**). Bcl-xL is a member of the Bcl-2 family of proteins, which regulate apoptosis by controlling mitochondrial outer membrane permeabilization^[29,30]^. It localizes at the outer mitochondrial membrane via a C-terminal transmembrane domain and functions as an anti-apoptotic factor by binding and sequestering proteins such as BAK and BAD via their Bcl-2 homology 3 (BH3) domains^[31,32]^. This well-characterized, high-affinity interaction makes it a reliable model for detecting heterodimeric protein-protein interactions. For imaging the interaction between Bcl-xL and BH3, we fused CFAST2 to a modified version of Bcl-xL in which the C-terminal trans membrane domain is replaced by the transmembrane domain of TOM20 (residues 1-34) to ensure precise localization at mitochondria (and avoid ER localization), while we fused NFAST2 to the BH3 domain of BAK. Co-expression of these constructs in cells led to the successful complementation of splitFAST2 at the mitochondria. The resolution enhancement achieved through STED microscopy enabled to demonstrate the specific localization of this interaction at the outer mitochondrial membrane, further underscoring the powerful combination of splitFAST2 and STED microscopy for sub-diffraction mapping of protein-protein interactions in whole, live cells.

To further highlight the versatility of splitFAST2 for sub-diffraction imaging, we expanded its use beyond PPI, focusing on proteins binding to adjacent structural unit. As model system, we fused the two halves of splitFAST2 to LifeAct (**Supplementary Fig. 7** and **Fig. 5**). This approach was designed to enable low-background imaging of F-actin as fluorescence is generated only when the two halves bind to adjacent actin monomers within F-actin filaments, while unbound proteins diffusing freely in cell remain non-fluorescent. We designed two fusions by tethering LifeAct at the C-terminus of NFAST2 or at the N-terminus of FKBP-CFAST2. Note that, as LifeAct and CFAST2 are two small peptides, we included FKBP in the LifeAct-FKBP-CFAST2 fusion to avoid proteolytic degradation. Confocal and STED microscopy of cells expressing these fusions revealed thin actin filaments with low background signal compared to control cells expressing the fusion LifeAct-pFAST containing full-length pFAST (**Fig. 5a,b**). This demonstrated the ability of splitFAST2 to enhance the signal-to-background ratio in this context. One interest of reducing background from diffusing proteins is to improve the quality and resolution of images. In STED microscopy, we could observe fine actin filament networks and bundles, such as those in microvilli and filopodia, achieving a full width at half-maximum of 90 nm compared to 300 nm in confocal microscopy (**Fig. 5c-f**).

**Figure 5.**
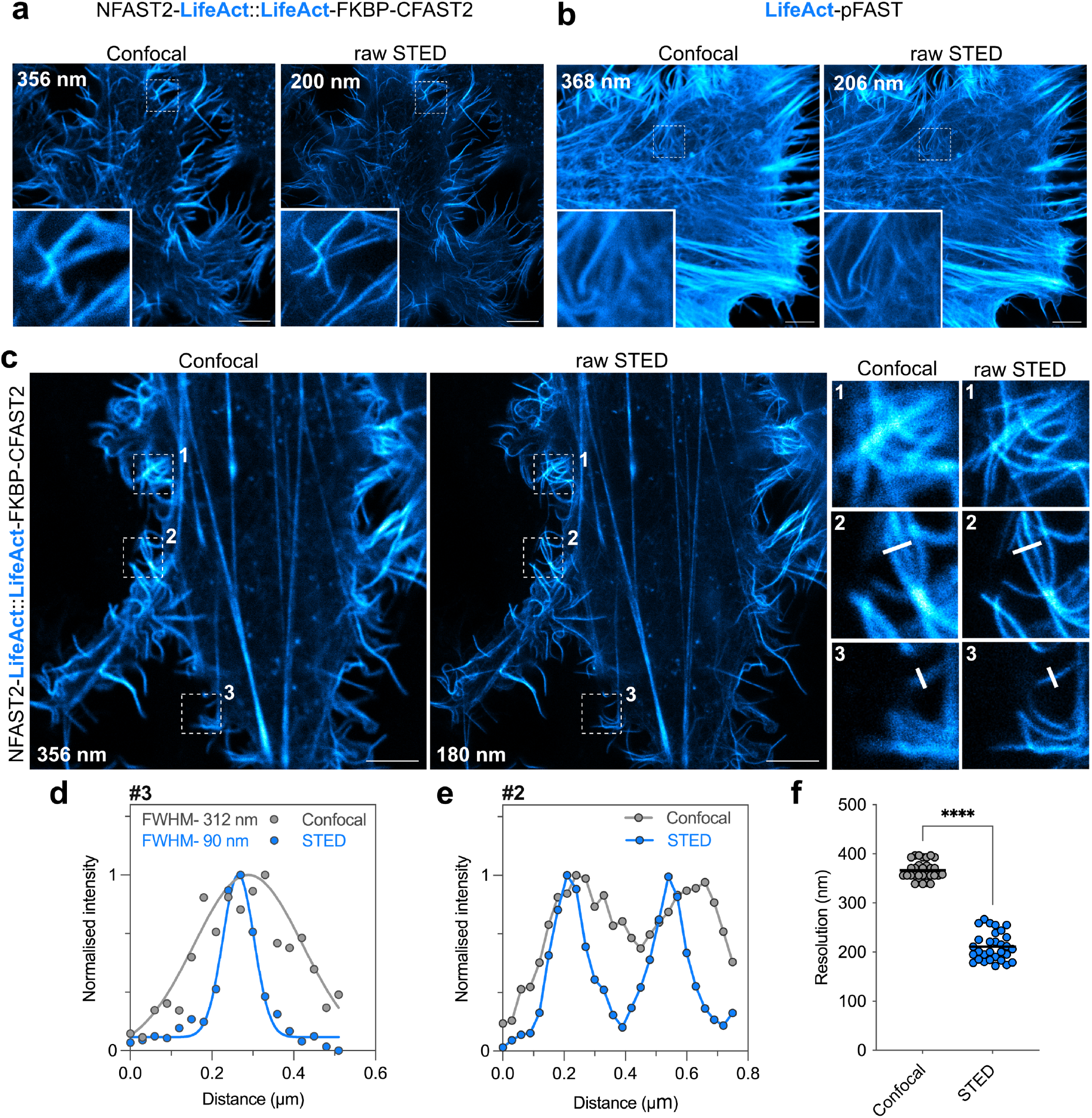
Live-cell subdiffraction imaging of filamentous actin with low-background splitFAST2-based fluorescent probes. **a**,**b** Confocal and raw STED micrographs of live HeLa cells expressing NFAST2-LifeAct and LifeAct-FKBP-CFAST2 (**a**) or LifeAct-pFAST (**b**) treated with 10 µM of HBR-3,5DOM. Global image resolution is given on the image. Scale bars, 5 µm. **c-f** Confocal and raw STED micrographs of live HeLa cells expressing NFAST2-LifeAct and LifeAct-FKBP-CFAST2 treated with 10 µM of HBR-3,5DOM. Three regions of interest were selected for close-up comparison. Micrographs are representative of n = 30 cells from three independent experiments. Global image resolution is given on the image. Scale bars, 5 µm. **d**,**e** Line profile across actin filaments from close-up were used to compare gain in resolution between confocal and STED. Graph shows raw data of line profile from confocal and STED images (points), in addition to the corresponding gaussian fit (line) in the case of (**d**). **f** Comparison of resolution of confocal and STED micrographs using image decorrelation analysis^[25]^ (n = 30 cells). See supplementary Table xxx for detailed acquisition parameters.

Expanding on these results, we adapted the approach for background-free imaging of microtubules. By fusing splitFAST2 fragments to MAP4, we engineered NFAST2-MAP4 and MAP4-CFAST2 constructs that only assemble when bound to adjacent tubulin monomers (**Supplementary Fig. 8**). The resulting probes facilitated high resolution STED imaging of microtubules in HBR-3,5DOM-treated cells, particularly in dense bundles (**Supplementary Fig. 9**). Beyond further demonstrating the suitability of splitFAST2 for sub-diffraction imaging of proximal proteins, this set of experiments underscore the high potential of splitFAST2 for the design of background-free imaging probes enabling enhanced contrast and resolution.

## CONCLUSION

In this study, we harnessed the power of the splitFAST2 technology in tandem with STED nanoscopy to visualize PPIs at subdiffraction resolution in whole living cells. By using a complementation-based reporter, fluorescence is triggered exclusively at sites where the two proteins interact, ensuring highly precise and unambiguous localization. This approach allowed us to map PPIs with high accuracy. Imaging PPI at subdiffraction resolution in whole, living cells is inherently challenging due to cell dynamics, which can degrade spatial resolution. In this context, scanning methods such as STED nanoscopy offer rapid imaging capabilities, making them particularly advantageous over SMLM approaches for subdiffraction imaging of PPI, even if the ultimate spatial resolution is somewhat lower. We demonstrated that combining splitFAST2 with STED enabled subdiffraction imaging of chemically inducible interactions, based on the FRB-FKBP-rapamycin system, or constitutive interactions such as the interaction between LifeAct and F-actin, the homodimerization of Lamin A/C or the interaction between Bcl-xL and the BH3 domain. The gain in resolution enabled to map the subcellular localization of these interactions with enhanced spatial resolution. In addition to facilitating the subdiffraction detection of PPI in whole living cells, our work demonstrates that splitFAST2 can be harnessed to create fluorescent probes with exceptionally low background for repetitive cellular structures such as F-actin and microtubules. This strategy is specifically designed so that fluorescence is only activated when both halves of splitFAST2 bind to adjacent monomers within filaments, effectively minimizing background signal from unbound proteins. This generalizable strategy opens the door for creating background-free probes that can be adapted to a wide array of cellular structures targeted by specific protein binders.

## ACKNOWLEDGMENTS

We thank the microscopy facility of the Institut de Biologie Paris Seine of Sorbonne University, and more particularly France Lam and Chloé Chaumeton for their assistance. We thank the ICM-QUANT quantitative Cellular and Molecular Imaging platform of the Paris Brain Institute. We thank Lydia Danglot from NeurImag Imaging core facility, part of the IPNP, Inserm 1266 unit and Université de Paris Cité for helping us with preliminary STED tests. This work has been supported by the Agence Nationale de la Recherche (ANR FluNanoTrack ANR-21-CE11-0010), the Institut Universitaire de France, and the Dynamic Imaging program of the Chan-Zuckerberg Initiative DAF (grant number 2023-321185), an advised fund of Silicon Valley Community.

## AUTHOR CONTRIBUTIONS

S.B. and A.G. designed the overall project and wrote the paper. S.B. and A.G. designed the experiments. S.B. performed the experiments. S.B. and A.G. analyzed the experiments.

