## Supporting Information for "Super-resolution live-cell mapping of protein-protein interactions using chemogenetic split reporters and STED microscopy"

**This pdf contains**

Supplementary Figures 1-9

Materials and Methods

Supplementary Tables 1-5

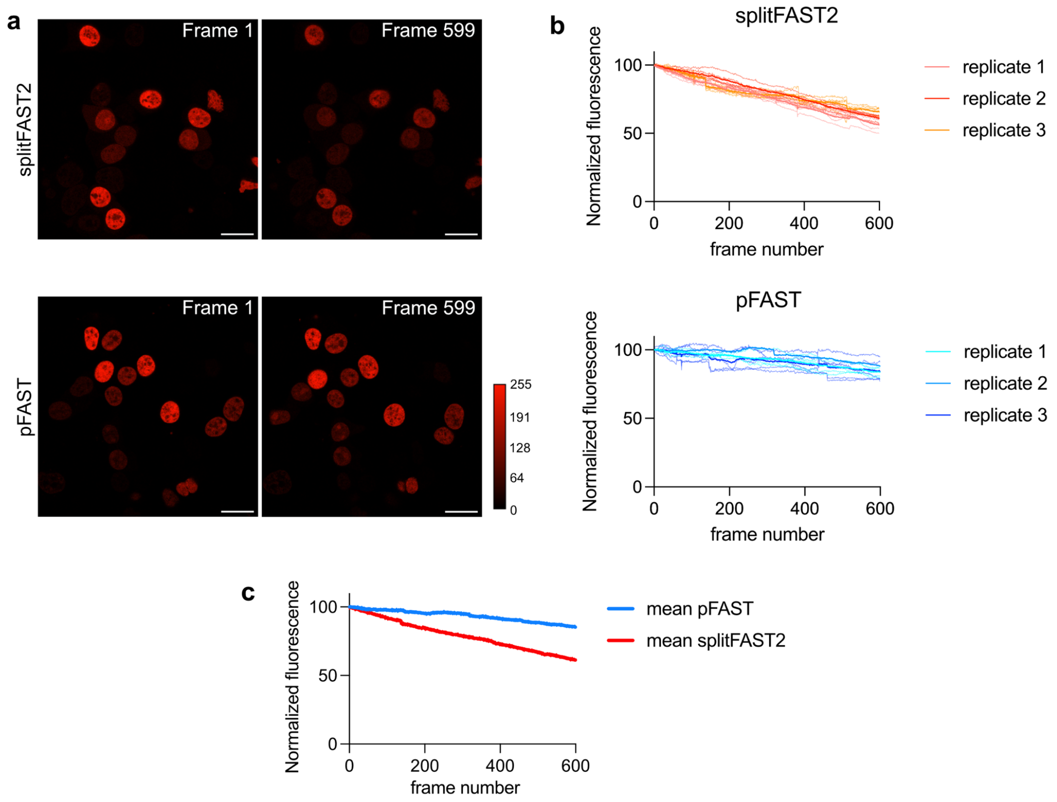


**Supplementary Figure 1. Photostability of splitFAST2 versus pFAST in confocal microscopy. a-c** HeLa cells expressing H2B-FRB-NFAST2 and FKBP-CFAST2 were treated with 100 nM of rapamycin and 10 µM HBR-3,5DOM and then imaged by confocal microscopy for 599 frames. Photostability is compared to that of H2B-pFAST imaged in the same conditions. **a** Representative micrographs. Scale bars 20 µm. See **Supplementary Table 1** for detailed acquisition parameters. **b** Fluorescence levels of 15 cells from three independent experiments over time. Each cell is color coded according to the independent experiment it came from. The thick lines correspond to the means of each replicate. **c** Average fluorescence of pFAST and splitFAST2 over time from experiments shown in **b**.

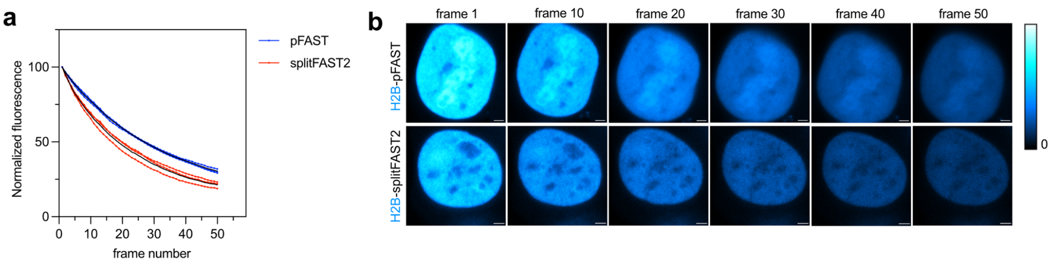


**Supplementary Figure 2. Photostability of splitFAST2 versus pFAST in STED microscopy.** HeLa cells expressing H2B-FRB-NFAST2 and FKBP-CFAST2 were treated with 100 nM of rapamycin and 10 µM HBR-3,5DOM and then imaged by STED microscopy for 50 frames. Photostability is compared to that of H2B-pFAST imaged in the same conditions. **a** Average fluorescence of pFAST and splitFAST2 (black lines) over time from three independent experiments. **b** Representative micrographs. Scale bars 2 µm. See **Supplementary Table 1** for detailed acquisition parameters.

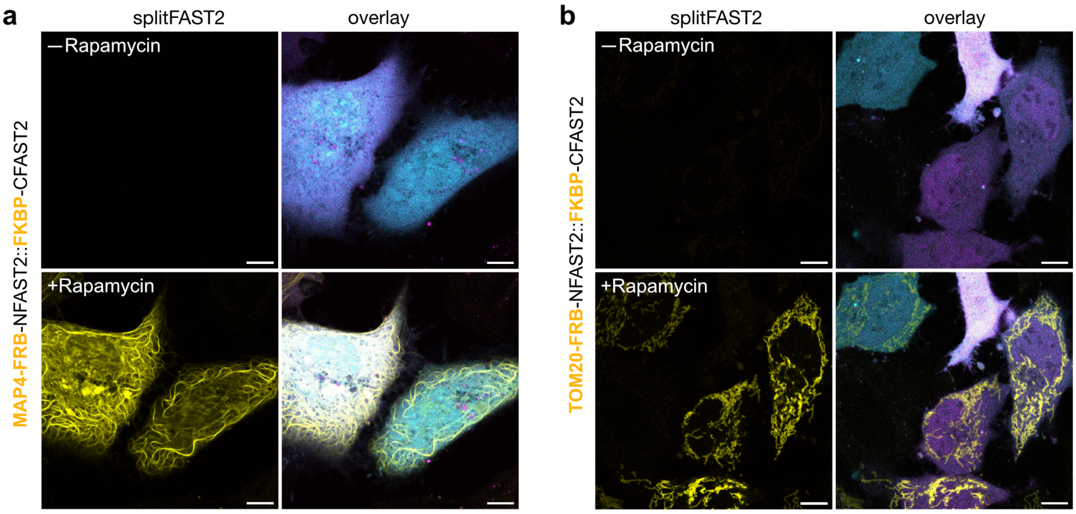


**Supplementary Figure 3. Live-cell imaging of chemically induced PPI using splitFAST2.** FRB-NFAST2 fusions and FKBP-CFAST2 were expressed in HeLa cells using bicistronic plasmids allowing the cytosolic expression of mTurquoise 2 and iRFP670 as transfection reporters. Cells were prelabeled with 10 μM HBR-3,5DOM, and imaged before and after treatment with 100 nM rapamycin. Representative images before and after rapamycin addition are shown. The overlay images contain the signal from splitFAST2 (yellow), mTuquoise2 (cyan) and iRFP670 (magenta). FRB-NFAST2 fusions used: (**a**) MAP4-FRB-NFAST2, (**b**) TOM20-FRB-NFAST2. Scale bars 10 μm. See **Supplementary Table 1** for detailed acquisition parameters.

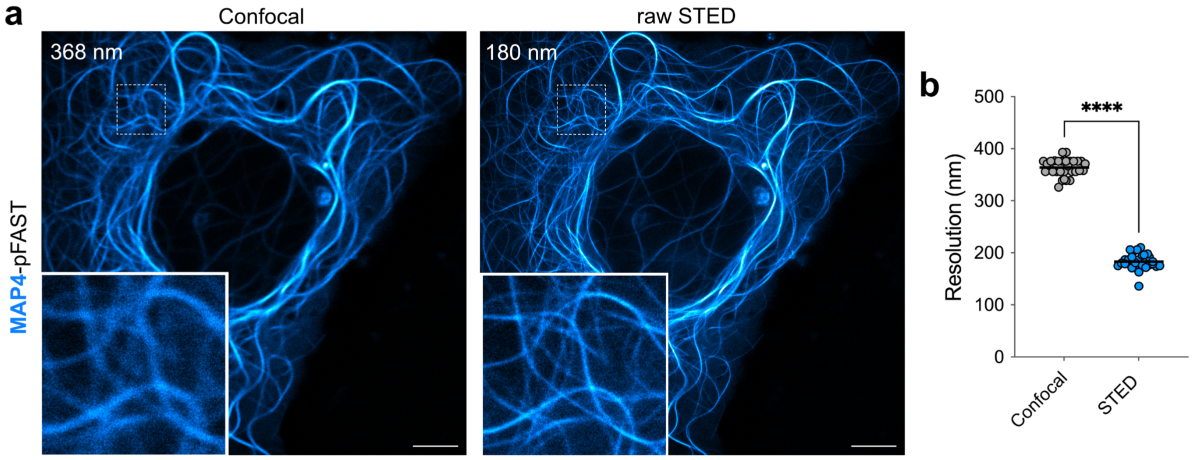


**Supplementary Figure 4. STED imaging of MAP4-pFAST. a** Confocal and raw STED micrographs of live HeLa cells expressing MAP4-pFAST treated with 10 µM of HBR-3,5DOM. Micrographs are representative of n = 30 cells from three independent experiments. Scale bars, 5 µm. See **Supplementary Table 1** for detailed acquisition parameters. **b** Comparison of resolution of confocal and STED micrographs using image decorrelation analysis[1] (n = 30 cells)

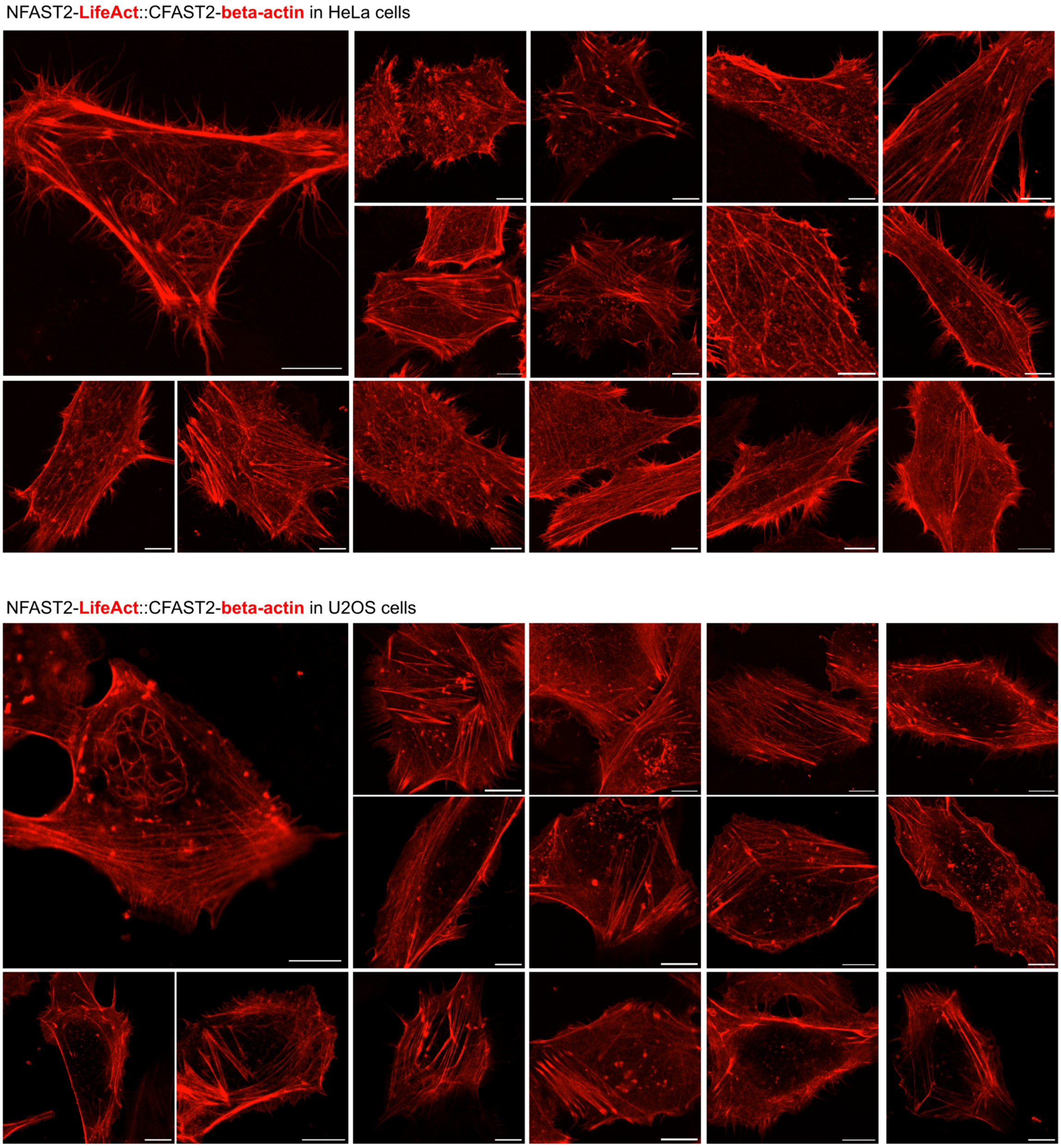


**Supplementary Figure 5. Live-cell confocal imaging of F-actin-LifeAct interaction.** Portfolio of Airyscan confocal micrographs of live HeLa and U2OS cells expressing NFAST2-LifeAct and CFAST2-β-actin pretreated with 10 μM of HBR-3,5DOM. Micrographs are representative of n = 30 cells from three independent experiments. Scale bars, 10 μm. See **Supplementary Table 1** for detailed acquisition parameters.

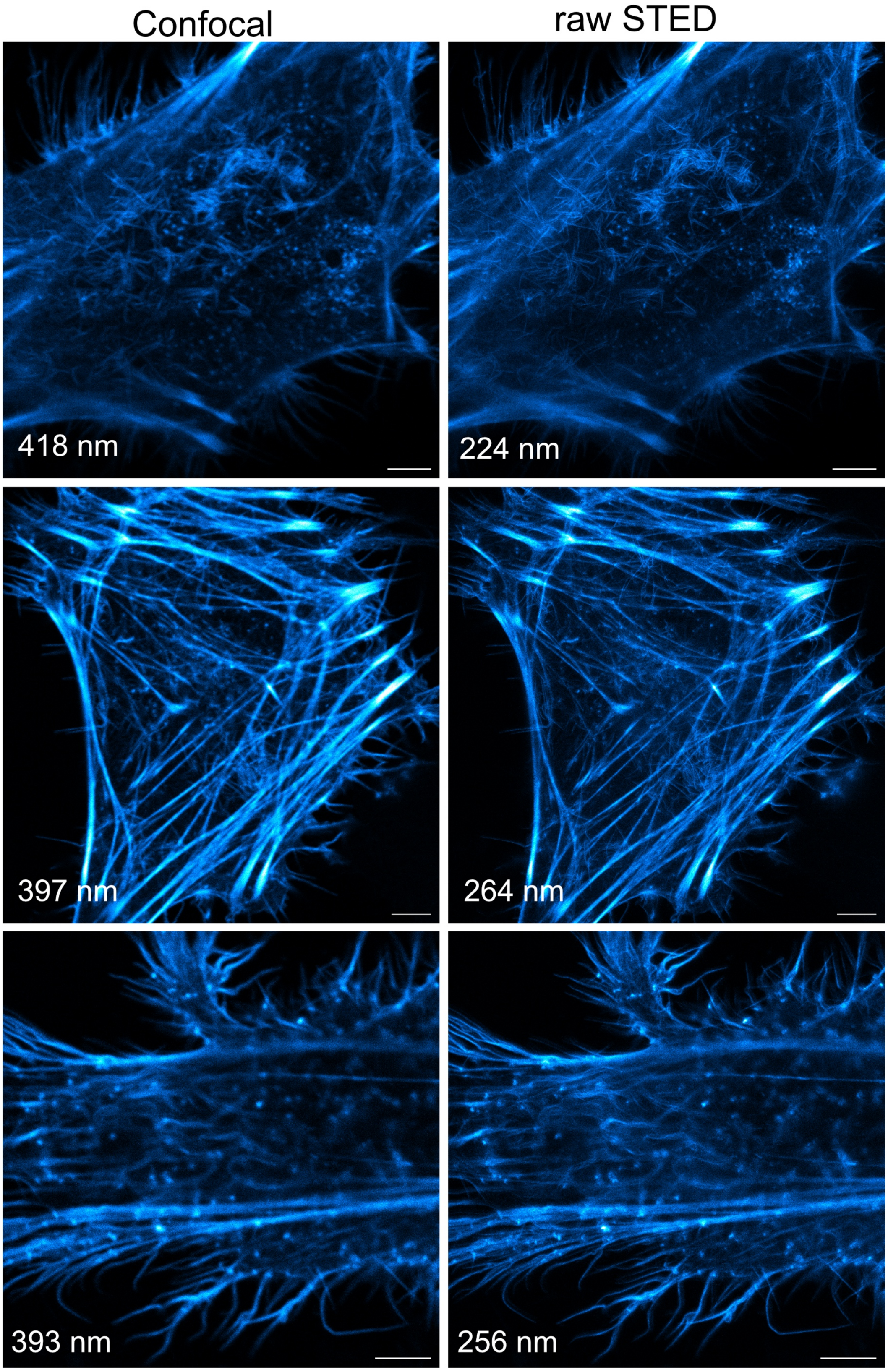


**Supplementary Figure 6. Live-cell STED imaging of F-actin-LifeAct interaction (1/6).** Portfolio of confocal and raw STED micrographs of live HeLa expressing NFAST2-LifeAct and CFAST2-β-actin pretreated with 10 μM of HBR-3,5DOM. Scale bars, 5 μm. See **Supplementary Table 1** for detailed acquisition parameters.

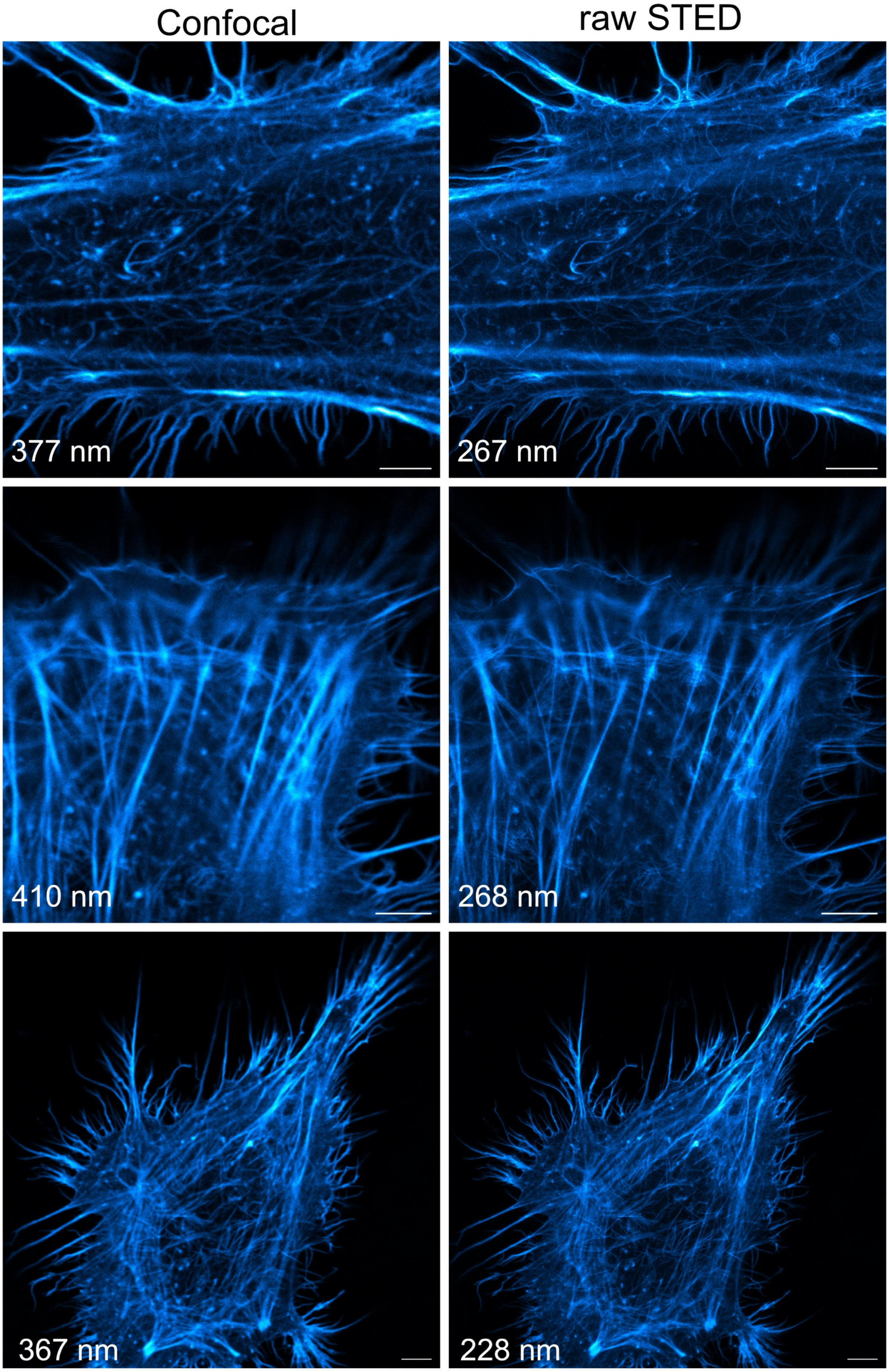


**Supplementary Figure 6. Live-cell STED imaging of F-actin-LifeAct interaction (2/6).** Portfolio of confocal and raw STED micrographs of live HeLa expressing NFAST2-LifeAct and CFAST2-β-actin pretreated with 10 µM of HBR-3,5DOM. Scale bars, 5 µm. See **Supplementary Table 1** for detailed acquisition parameters.

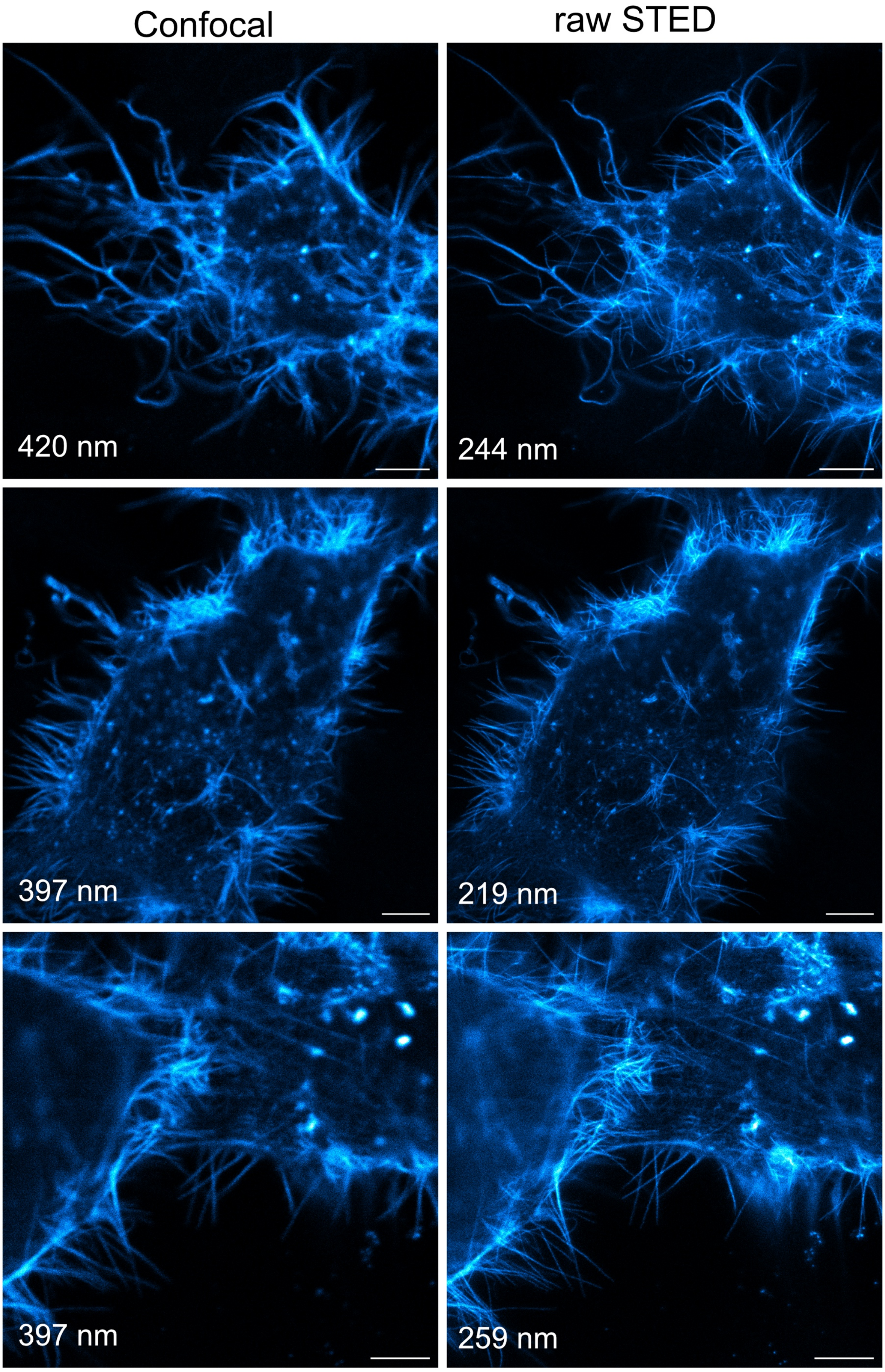


**Supplementary Figure 6. Live-cell STED imaging of F-actin-LifeAct interaction (3/6).** Portfolio of confocal and raw STED micrographs of live HeLa expressing NFAST2-LifeAct and CFAST2-β-actin pretreated with 10 μM of HBR-3,5DOM. Scale bars, 5 μm. See **Supplementary Table 1** for detailed acquisition parameters.

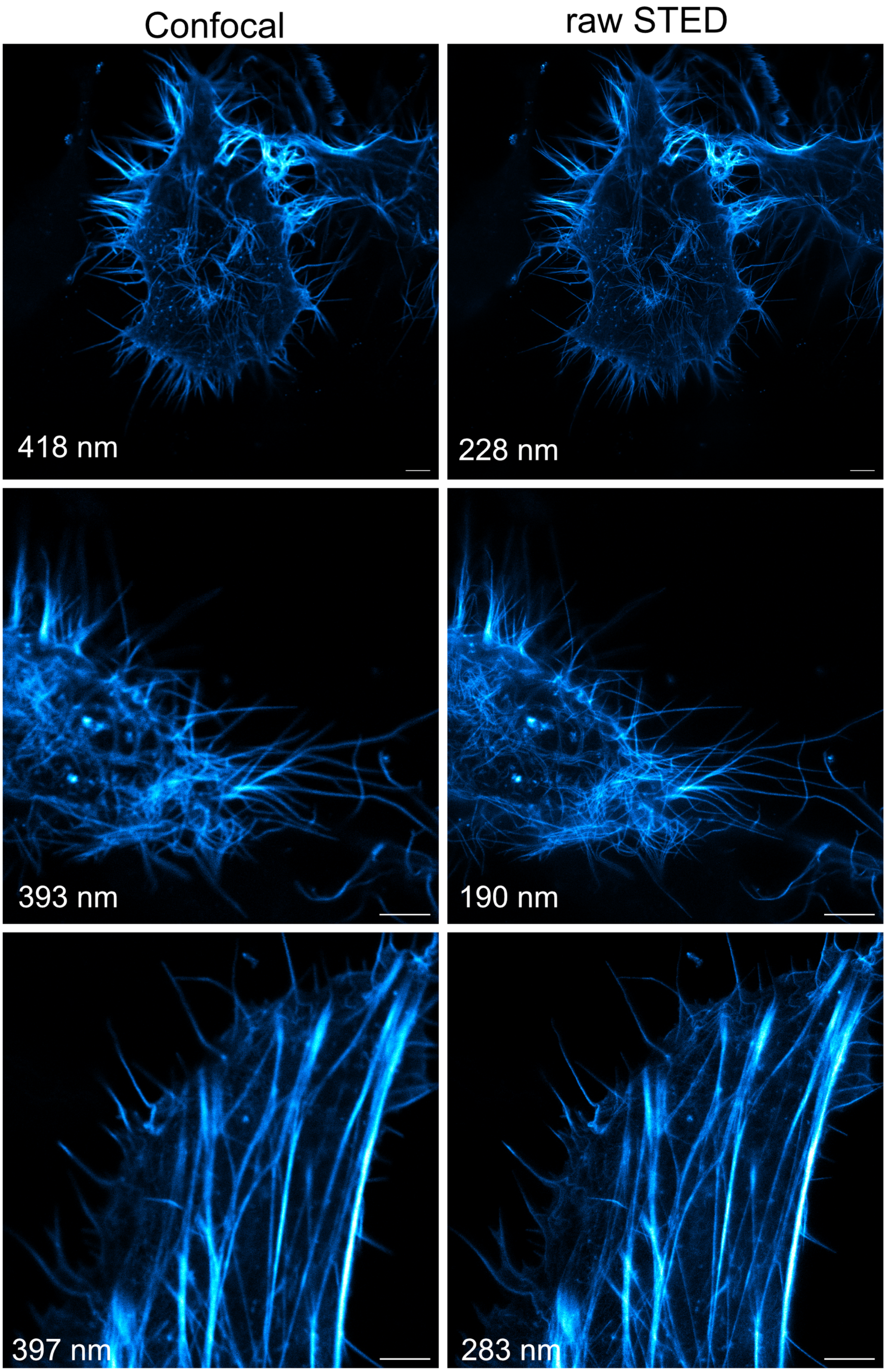


**Supplementary Figure 6. Live-cell STED imaging of F-actin-LifeAct interaction (4/6).** Portfolio of confocal and raw STED micrographs of live HeLa expressing NFAST2-LifeAct and CFAST2-β-actin pretreated with 10 μM of HBR-3,5DOM. Scale bars, 5 μm. See **Supplementary Table 1** for detailed acquisition parameters.

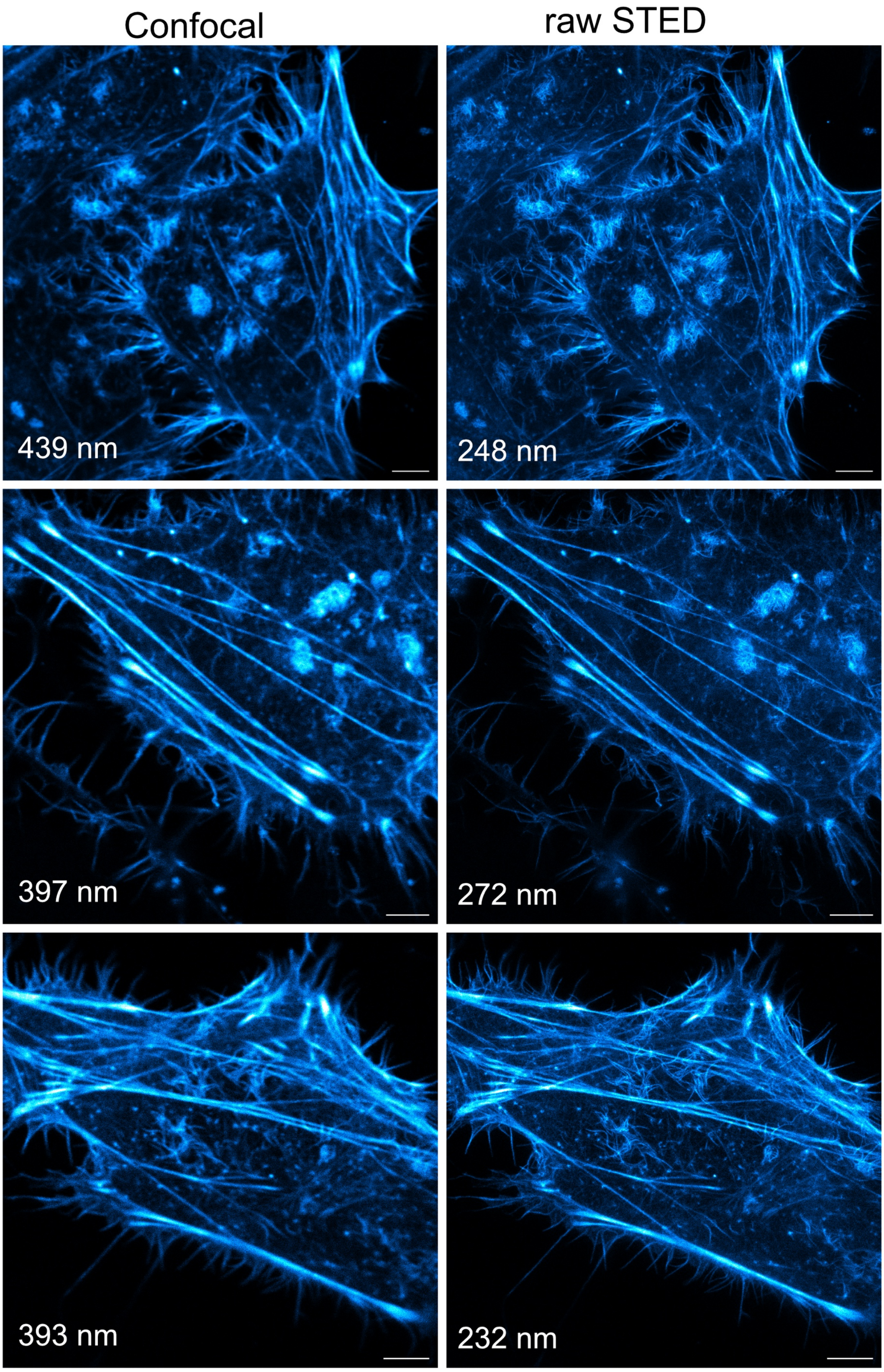


**Supplementary Figure 6. Live-cell STED imaging of F-actin-LifeAct interaction (5/6).** Portfolio of confocal and raw STED micrographs of live HeLa expressing NFAST2-LifeAct and CFAST2-β-actin pretreated with 10 µM of HBR-3,5DOM. Scale bars, 5 µm. See **Supplementary Table 1** for detailed acquisition parameters.

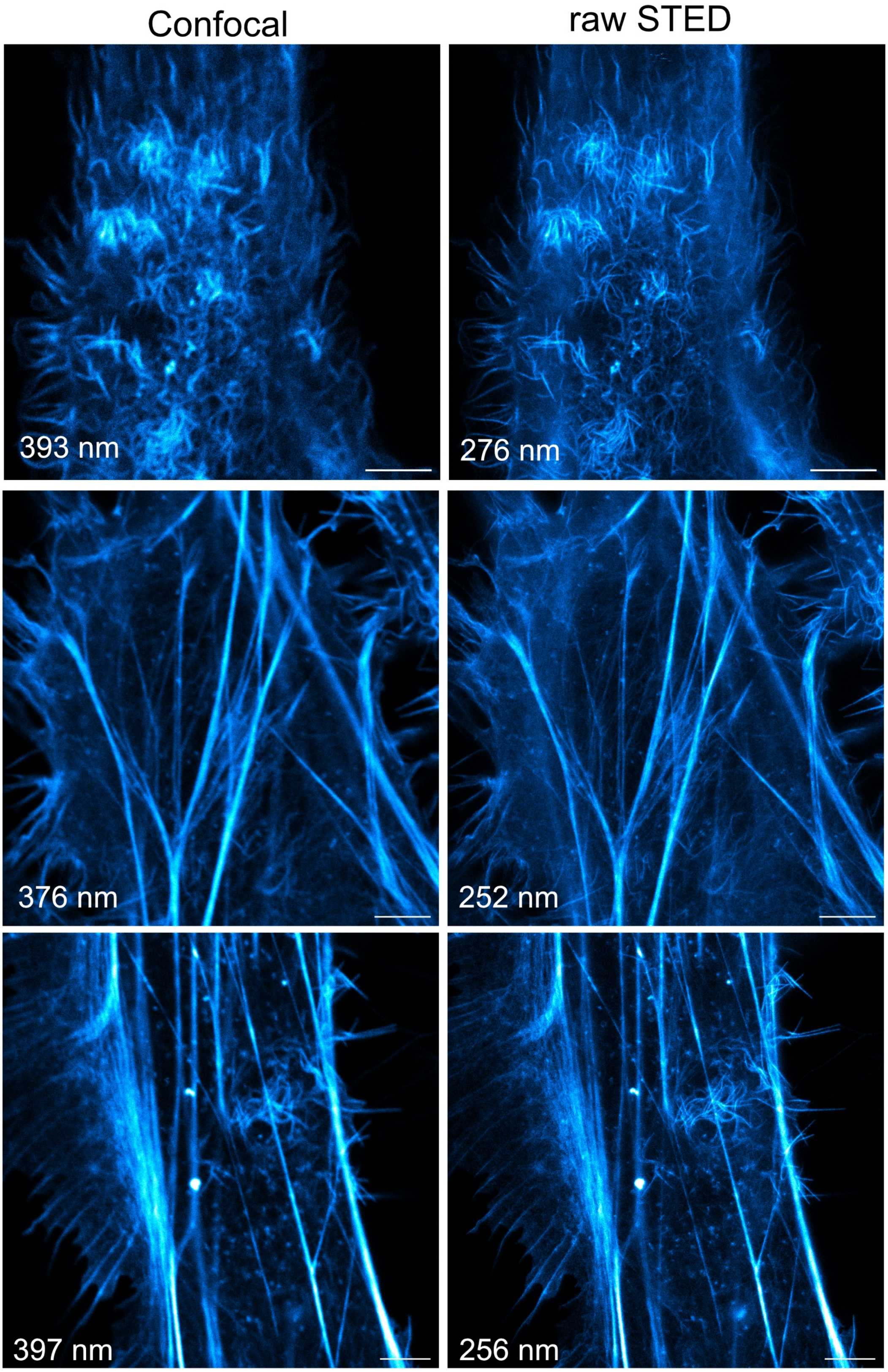


**Supplementary Figure 6. Live-cell STED imaging of F-actin-LifeAct interaction (6/6).** Portfolio of confocal and raw STED micrographs of live HeLa expressing NFAST2-LifeAct and CFAST2-β-actin pretreated with 10 μM of HBR-3,5DOM. Scale bars, 5 μm. See **Supplementary Table 1** for detailed acquisition parameters.

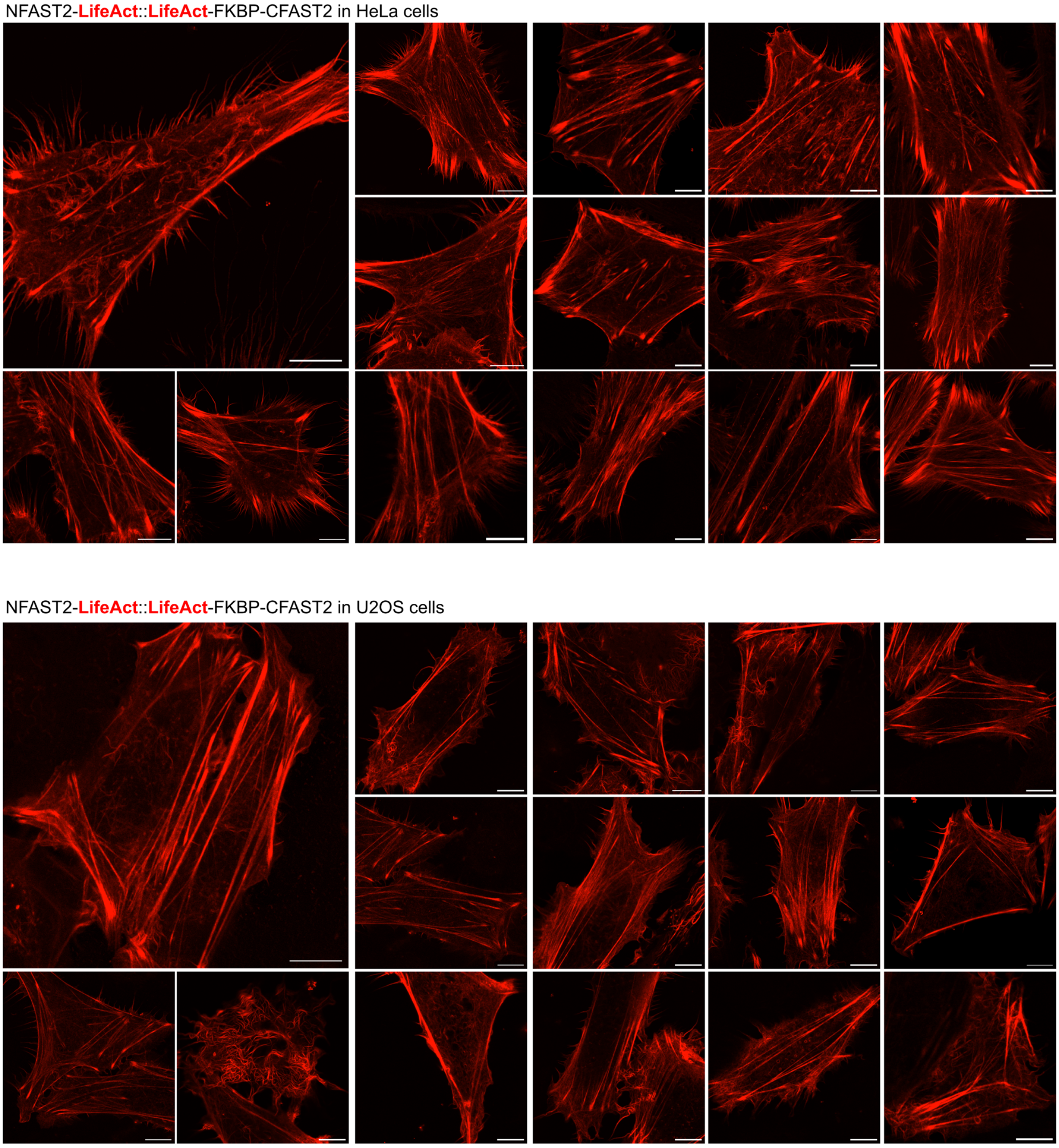


**Supplementary Figure 7. Live-cell confocal imaging of F-actin with low-background splitFAST2-based fluorescent probes.** Portfolio of Airyscan confocal micrographs of live HeLa and U2OS cells expressing NFAST2-LifeAct and LifeAct-FKBP-CFAST2-LifeAct pretreated with 10 µM of HBR-3,5DOM. Micrographs are representative of n = 30 cells from three independent experiments. Scale bars, 10 µm. See **Supplementary Table 1** for detailed acquisition parameters.

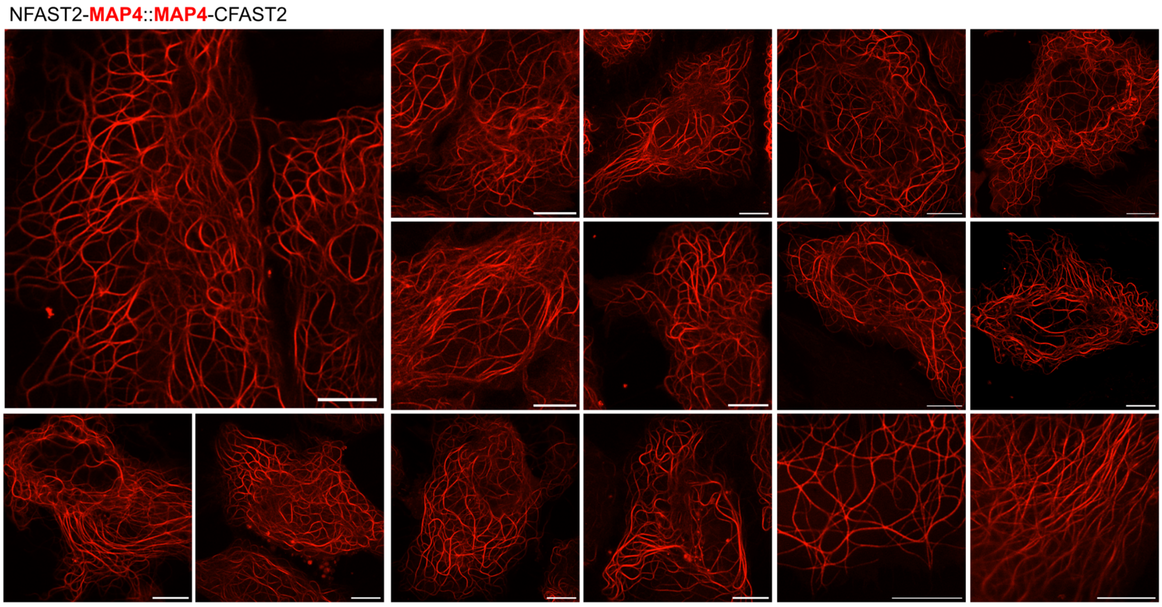


**Supplementary Figure 8. Live-cell confocal imaging of microtubules with low-background splitFAST2-based fluorescent probes.** Portfolio of Airyscan confocal micrographs of live HeLa expressing NFAST2-MAP4 and MAP4-CFAST2-LifeAct pretreated with 10 µM of HBR-3,5DOM. Micrographs are representative of n = 30 cells from three independent experiments. Scale bars, 10 µm. See **Supplementary Table 1** for detailed acquisition parameters.

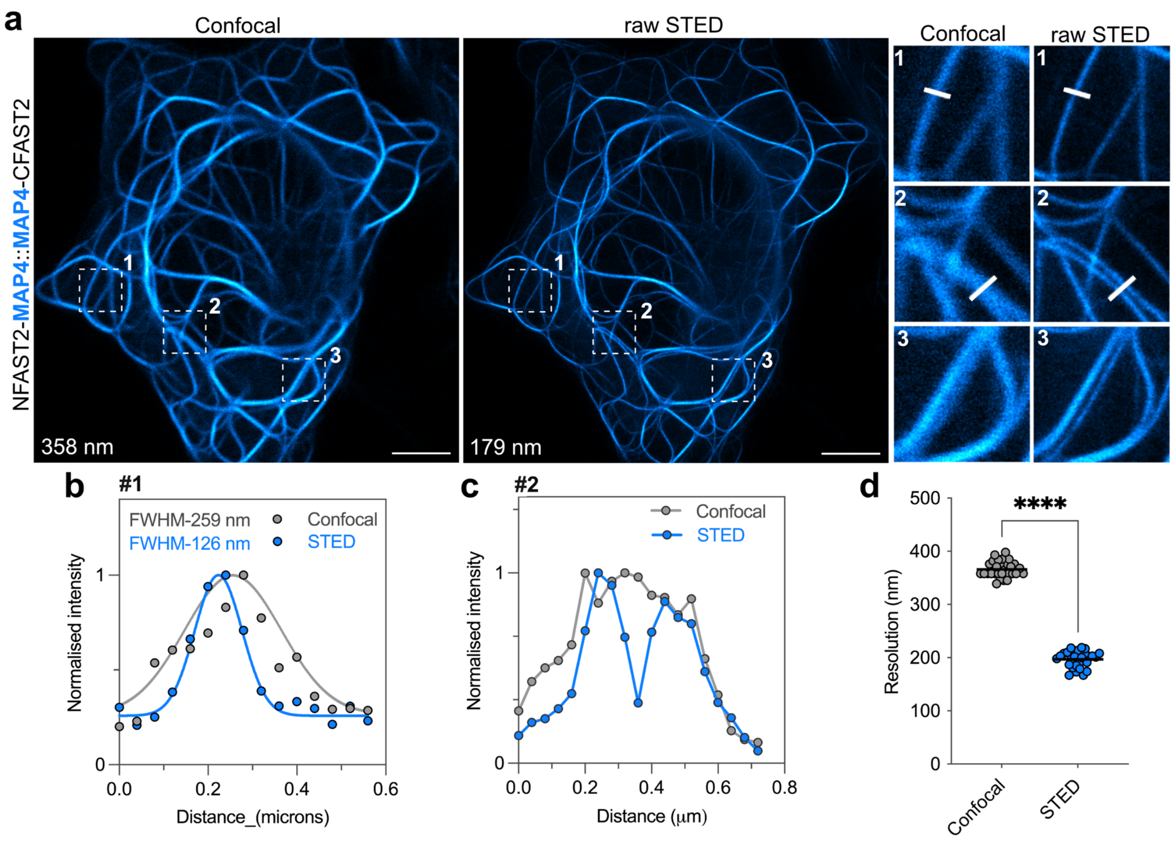


**Supplementary Figure 9. Live-cell subdiffraction imaging of microtubules with low-background splitFAST2-based fluorescent probes. a** Confocal and raw STED micrographs of live HeLa cells expressing NFAST2-MAP4 and MAP4-CFAST2 treated with 10 μM of HBR-3,5DOM. Three regions of interest were selected for close-up comparison. Micrographs are representative of n = 30 cells from three independent experiments. Scale bars, 5 μm. See **Supplementary Table 1** for detailed acquisition parameters. **b,c** Line profile across microtubules from close-up were used to compare gain in resolution between confocal and STED. Graph shows raw data of line profile from confocal and STED images (points), in addition to the corresponding gaussian fit (line) in the case of (**b**). **d** Comparison of resolution of confocal and STED micrographs using image decorrelation analysis[1] (n = 30 cells).

### Supplementary Table 1. Imaging settings

| Figure | Panel | Fluorescent reporter | Excitation (nm) | Emission (nm) | Imaging setup | Comments |
|---|---|---|---|---|---|---|
| **Fig. 2** | **a** | splitFAST2 HBR-3,5DOM | 561 | Abberior STAR Orange | Abberior facility Line STED 775 nm depletion laser 60 NA1.42 × oil | pixel size 30 nm Dwell time 5 µs Line averaging × 6 STED power 15 % |
| | **e** | splitFAST2 HBR-3,5DOM | 561 | Abberior STAR Orange | Abberior facility Line STED 775 nm depletion laser 60 NA1.42 × oil | pixel size 30 nm Dwell time 5 µs Line averaging × 6 STED power 18 % |
| **Fig. 3** | **a** | splitFAST2 HBR-3,5DOM | 561 | Abberior STAR Orange | Abberior facility Line STED 775 nm depletion laser 60 NA1.42 × oil | pixel size 30 nm Dwell time 5 µs Line averaging × 6 STED power 20 % |
| | **e** | splitFAST2 HBR-3,5DOM | 561 | Abberior STAR Orange | Abberior facility Line STED 775 nm depletion laser 60 NA1.42 × oil | pixel size 30 nm Dwell time 5 µs Line averaging × 6 STED power 20 % |
| Fig. 4 | **a** | splitFAST2 HBR-3,5DOM | 561 | Abberior STAR Orange | Abberior facility Line STED 775 nm depletion laser 60 NA1.42 × oil | pixel size 30 nm Dwell time 5 µs Line averaging × 6 STED power 10 – 15 % |
| | **e** | splitFAST2 HBR-3,5DOM | 561 | Abberior STAR Orange | Abberior facility Line STED 775 nm depletion laser 60 NA1.42 × oil | pixel size 30 nm Dwell time 5 µs Line averaging × 6 STED power 20 % |
| Fig. 5 | **a** | splitFAST2 HBR-3,5DOM | 561 | Abberior STAR Orange | Abberior facility Line STED 775 nm depletion laser 60 NA1.42 × oil | pixel size 30 nm Dwell time 5 µs Line averaging × 6 STED power 20 % |

| | | | | | | |
|---|---|---|---|---|---|---|
| | b | pFAST HBR-3,5DOM | 561 | Abberior STAR Orange | Abberior facility Line STED 775 nm depletion laser 60 NA1.42 × oil | pixel size 30 nm Dwell time 5 μs Line averaging × 6 STED power 20 % |
| | c | splitFAST2 HBR-3,5DOM | 561 | Abberior STAR Orange | Abberior facility Line STED 775 nm depletion laser 60 NA1.42 × oil | pixel size 30 nm Dwell time 5 μs Line averaging × 6 STED power 20 % |
| SI Fig. 1 | a | splitFAST2 HBR-3,5DOM & pFAST HBR3,5DOM | 561 | 570-650 | LSM980 inverted 63 × oil | 1 frame / 2 s Power: 2% 1 frame / 2 s |
| SI Fig. 2 | | splitFAST2 HBR-3,5DOM & pFAST HBR-3,5DOM | 561 | Abberior STAR Orange | Abberior facility Line STED 775 nm depletion laser 60 NA1.42 × oil | pixel size 30 nm Dwell time 5 μs Line averaging × 3 STED power 10 % 1 frame /10 s |
| SI Fig. 3 | a,b | splitFAST2 HBR-3,5DOM<br>mTurquoise2<br>iRFP670 | 561<br>445<br>639 | 570-650<br>470-520<br>650-700 | LSM980 inverted 63 × oil | 1 frame / 2 s |
| SI Fig. 4 | a | splitFAST2 HBR-3,5DOM | 561 | Abberior STAR Orange | Abberior facility Line STED 775 nm depletion laser 60 NA1.42 × oil | pixel size 30 nm Dwell time 5 μs Line averaging × 6 STED power 20 % |
| SI Fig. 5 | | splitFAST2 HBR-3,5DOM | 561 | 570-650 | LSM980 inverted 63 × oil | Airyscan mode |
| SI Fig. 6 | a | splitFAST2 HBR-3,5DOM | 561 | Abberior STAR Orange | Abberior facility Line STED 775 nm depletion laser 60 NA1.42 × oil | pixel size 30 nm Dwell time 5 μs Line averaging × 6 STED power 20% |

| | | | | | |
|---|---|---|---|---|---|
| SI Fig. 7 | splitFAST2<br>HBR-3,5DOM | 561 | 570-650 | LSM980 inverted<br>63 × oil | Airyscan mode |
| SI Fig. 8 | splitFAST2<br>HBR-3,5DOM | 561 | 570-650 | LSM980 inverted<br>63 × oil | Airyscan mode |
| SI Fig. 9 | splitFAST2<br>HBR-3,5DOM | 561 | Abberior<br>STAR Orange | Abberior facility Line STED<br>775 nm depletion laser<br>60 NA1.42 × oil | pixel size<br>30 nm<br>Dwell time 5 μs<br>Line averaging<br>× 6<br>STED power<br>15% |

### Materials and methods

#### General

Synthetic oligonucleotides used for cloning were purchased from Integrated DNA Technology. PCR reactions were performed with Q5 polymerase (New England Biolabs) following manufacturer guidelines. PCR products and restriction digest fragments were loaded to 1-2% agarose gel for 45 min at 100 V in TAE (Tris, Acetate, EDTA) buffer and purified using QIAquick Gel Extraction Kit PCR purification kit (Qiagen). Cloning was performed using restriction digest and Gibson assembly methods. Restriction enzymes were purchased by NEB or Thermofisher scientific. Isothermal assembly (Gibson assembly) was performed using home-made isothermal reaction buffer prepared according to previously described protocols[2]. Hybridisation of short oligos were performed using a ramp down method in a thermocycler starting at 95°C reducing 1°C per cycle for 70 cycles. Constructs were ligated using T4 ligase following manufacturer guidelines (NEB). All products were transformed in DH10 beta (NEB) *E. coli* strains using either electroporation or chemically competent methods as described by manufacturers. Small-scale isolation of plasmid DNA was done using QIAprep miniprep kit (Qiagen) from 3 mL of overnight culture supplemented with appropriate antibiotics. Large-scale isolation of plasmid DNA was done using the QIAprep maxiprep kit (Qiagen) from 150 mL of overnight culture supplemented with appropriate antibiotics. All constructs were cloned into a common vector with a pBR322 origin of replication site and a CMV promoter sequence. All plasmid sequences were confirmed by Sanger sequencing with appropriate sequencing primers or whole plasmid sequencing (Eurofins genomics).

#### Mammalian cell culture

All reagents were purchased from Thermofisher scientific unless stated otherwise. HEK 293T cells (ATCC CRL-3216) were cultured in Dulbecco's modified Eagle's medium (DMEM) supplemented with high Glucose, L-glutamine, Phenol Red, Sodium Pyruvate and 10% (vol/vol) fetal calf serum, at 37 °C in a 5% $CO_2$ atmosphere. HeLa cells (ATCC CCL-2) were cultured in modified Eagle's medium (MEM) supplemented with phenol red, 1 × non-essential amino acids, 1× sodium pyruvate, and 10% (vol/vol) fetal calf serum at 37 °C in a 5% $CO_2$ atmosphere. U-2 OS (HTB-96) cells were cultured in McCoy media supplemented with 10% (vol/vol) fetal calf serum at 37 °C in a 5% $CO_2$ atmosphere. For imaging, cells were seeded in μDish 35 mm glass bottom dish (Ibidi) coated with poly-L-lysine. Cells were seeded 18 to 24 h prior to transfection at a concentration of 120,000 HEK 293T cells and 60,000 HeLa and U-

2 OS per dish. Cells were transiently transfected with 1 μg total plasmid DNA. Transfection was achieved using Genejuice (Merck) and reduced serum media OptiMEM according to the manufacturer's protocol 24 h prior to microscopy imaging.

**Fluorescence microscopy**

Confocal micrographs of mammalian cells were acquired on a Zeiss LSM 980 Laser Scanning Microscope equipped with a plan apochromat 63×/1.4 NA oil immersion objective. Stimulated emission depletion (STED) and confocal micrographs were acquired on an Abberior facility line microscope equipped with U-plan X apochromat 60×/1.42 NA oil immersion objective. Live cells were washed twice with DMEM media (without serum and phenol red) immediately prior to imaging. To the same DMEM media, the fluorogens were diluted at the indicated concentration and added to the cells. Cells were imaged directly without washing. ZEN software (for confocal imaging) and Imspector software (for STED imaging) were used to collect the data. The images were analysed with Fiji (Image J) and data processed using GraphPad Prism 7.

**Plasmid construction**

The plasmids used in these studies allowed the expression of proteins in mammalian cells. Expression is under the control of a CMV promoter.

Plasmids pAG657[3], pAG580[4] were previously described. Plasmid pAG1356 encoding MAP4-FRB-NFAST2-IRES2-mTurquoise2 was constructed by Gibson assembly from plasmids pAG573 (FRB-NFAST2–IRES2–mTurquoise2)[4] and pAG665 (MAP4-pFAST)[3]. The MAP4 insert was amplified using primers ag2194 and ag2197 with overlapping sequences containing BglII and SpeI restriction sites. The remaining vector was amplified from pAG573 in two fragments: fragment 1 with primers ag2195 and ag2088, incorporating a GGGSSSGGG linker, and fragment 2 with primers ag2196 and ag2089. All fragments were treated with DpnI, gel purified and assembled at 50°C for 1 hour in a thermocycler.

Plasmids pAG1359 (TOM20-FRB-NFAST2-*IRES2*-MT2) and pAG1965 (H2B-FRB-NFAST2-*IRES2*-MT2) were generated from the pAG1356 vector encoding MAP4-FRB-NFAST2-*IRES2*-mTurquoise2. For pAG1359, the sequence encoding TOM20 was amplified by PCR from Addgene plasmid #171461 using oligos ag2202 and ag2203 with BglII and SpeI sites. For pAG1965, the sequence encoding H2B was amplified from pAG657[3] using oligos ag3841 and ag3842, all with BglII and SpeI sites. The vector pAG1356 was digested with BglII and SpeI restriction enzymes to remove the MAP4 insert, and the linearised backbone was dephosphorylated with CIP. PCR products for TOM20 and H2B were digested with BglII and SpeI and purified.

The vector backbone of plasmids encoding NFAST2-*IRES2*-MT2 and CFAST2-*IRES2*-iRFP670 were first modified from plasmids pAG892 NFAST2-H3-*IRES2*-mTurquoise2[4] and pAG895 HP1-CFAST2-*IRES2*-iRFP670[4] by Gibson assembly. pAG892 was sequentially modified to introduce BamHI and AgeI sites flanking the gene of interest using oligos ag1948 and ag1950. The vector backbone was then altered by site-directed mutagenesis to remove an additional BamHI site using oligos ag2368 and ag2370, and the remaining vector was amplified with oligos ag2366 and ag2367. The resulting plasmid, pAG1457, served as the template for constructs containing the NFAST2 sequence. Vector pAG895 was similarly modified using oligos ag1935 and ag1952. To remove an NheI site in the original iRFP670 sequence, site-directed mutagenesis was performed with oligos ag1953 and ag1940, and the remaining vector was amplified using oligos ag1937 and ag1938. Further modification of the *IRES2* sequence was carried out with oligos ag2368 and ag2370. The final plasmid, pAG1428, was used as the template for constructs containing CFAST2. All fragments were treated with DpnI, gel purified and assembled at 50°C for 1 hour in a thermocycler.

To generate pAG1520 encoding NFAST2-LifeAct-IRES2-MT2, oligos ag2518 and ag2519 encoding the LifeAct peptide were hybridized with BamHI and AgeI overhangs using the ramp-down method, then ligated into BamHI/AgeI-digested pAG1457. pAG1522 encoding LifeAct-FKBP-CFAST2-*IRES2*-iRFP670 was generated by Gibson assembly using pAG1520 and pAG580 as templates. The FKBP sequence was amplified from pAG580 with oligos ag2563 and ag2089, and LifeAct was amplified from pAG1520 using oligos ag1858 and ag2562.The remaining vector sequence was amplified from pAG580 using oligos ag2561 and ag2088. Fragments were treated with DpnI, gel purified and assembled at 50°C for 1 hour in a thermocycler. pAG1754 encoding CFAST2-βActin-*IRES2*-iRFP670 was generated sequentially from pAG1457. The NFAST2 sequence was replaced with CFAST2 by digestion with BglII and BspEI, using hybridised oligos ag2522 and ag2523 to generate the CFAST2 fragment with sticky ends. The β-Actin coding sequence was ordered from Integrated DNA Technologies (IDT) and ligated into the CFAST2-BamHI-AgeI-*IRES2*-MT2 vector between BamHI and AgeI to produce pAG1528. The MT2 sequence was then removed by EcoRI and NotI digestion and replaced with iRFP670, excised from pAG1522 using the same enzymes. To generate pAG1530 (NFAST2-MAP4-*IRES2*-MT2) and pAG1531 (MAP4-CFAST2-*IRES2*-iRFP670), the MAP4 coding sequence was PCR-amplified from plasmid pAG665. BamHI and AgeI site mutations were introduced via Gibson assembly using oligo pairs ag2516/ag2089 and ag2517/ag2088 (BamHI) and ag2674/ag2089 and ag2675/ag2088 (AgeI). For pAG1530, MAP4 was PCR-amplified with a 5' BamHI site and a 3' stop codon plus AgeI site (oligos ag2676/ag2560), digested and ligated into a BamHI/AgeI-digested NFAST2-IRES2-MT2 vector. For pAG1531, MAP4 was amplified with a 5' NheI and 3' BamHI site (oligos ag2557/ag2558), digested, purified and ligated into a NheI/BamHI-digested CFAST2-IRES2-iRFP670. To generate pAG1936 encoding NFAST2-LaminA/C-*IRES2*-MT2, the DNA fragment encoding Lamin A/C (Addgene 55068) was amplified by PCR using oligos to modify the existing restriction site AgeI with oligo pairs ag3732/ag3733 and ag3734/ag3819. Oligos pairs ag3735/ag3818 and ag3736/ag3737, were used to amplify the plasmid backbone from pAG1530. Fragments were treated with DpnI, gel purified and assembled at 50°C for 1 h in a thermocycler. To generate pAG1976 encoding CFAST2-LaminA/C-*IRES2*-iRFP670, both Lamin A/C insert from pAG1936 and the plasmid backbone from pAG1754 were digested with BamHI and AgeI, the selected fragments were purified and ligated. To generate pAG1939 encoding NFAST2-BH3domain-*IRES2*-MT2, long oligos ag3758 and ag3759 encoding the BH3 domain of BAK1 (BCL2 Antagonist/Killer 1) with sticky ends for BamHI and AgeI restriction sites and the N and C terminal respectively, were hybridised using standard protocol. Fragments were ligated with previous digested vector containing NFAST2-BamHI-AgeI-IRES2-MT2. To generate pAG1967 encoding TOM20-Bcl-xL (Δ transmembrane domain)-CFAST2-*IRES2*-iRFP670, the TOM20 sequence was PCR-amplified from pAG1359 using oligos ag3762 and ag3765. The Bcl-xL coding sequence, excluding the C-terminal transmembrane domain, was ordered from IDT and amplified with oligos ag3764 and ag3767. The remaining vector was amplified from pAG1531 using oligo pairs ag3766/ag3735 and from pAG1359 using ag3736/ag3723. All fragments were treated with DpnI, gel purified and assembled at 50°C for 1 hour in a thermocycler.

**Mammalian cell transfection and analysis**

Mammalian cells were co-transfected with high purity DNA at a concentration of 1 µg of total plasmid DNA. For chemically induced interactions, ratios of 1-part pAG580 to 4-parts pAG1356, pAG1359 and pAG1965 were required for optimal interaction using transfection method described above. Positively transfected cells were first selected using the transfection reporter mTurquoise2 or iRFP670 placed after the *IRES2* sequence. To induce the interaction between rapamycin-binding protein (FKBP) and the FKBP-rapamycin binding domain (FRB), rapamycin was added into dishes to a final concentration of 100 nM. Cells were labelled with 10 µM HBR-3,5DOM. For constitutive interactions, ratios of 1-part NFAST2 to 1-part CFAST2 were required for optimal interaction using transfection method described above. Transfection efficiency cells was determined using the transfection reporter placed after the *IRES2* sequence, i.e. mTurquoise2 and iRFP670. Cells were labelled with 10 μM HBR-3,5DOM to monitor complementation efficiency of splitFAST2 fragments.

**Statistics & Reproducibility**

No sample size calculations were performed. When relevant, the sample size (n) is provided in the corresponding figure captions. Sample sizes were chosen to support meaningful conclusions. No data were excluded. The number of replicates for each individual experiments is indicated in the figure legends. All attempts at replication were successful. The experiments were not randomized. The Investigators were not blinded to allocation during experiments and outcome assessment.

**Supplementary Table 1. Plasmids**

| Plasmid | Resistance | Expression | Description |
|---|---|---|---|
| pAG1356 | kan | mammalian | CMV-MAP4-FRB-NFAST2-*IRES2*-MT2 |
| pAG1359 | kan | mammalian | CMV-TOM20-FRB-NFAST2-*IRES2*-MT2 |
| pAG1965 | kan | mammalian | CMV-H2B-FRB-NFAST2-*IRES2*-MT2 |
| pAG580 | kan | mammalian | CMV-FKBP-CFAST2-*IRES2*-iRFP670 |
| pAG657 | kan | mammalian | CMV-H2B-pFAST |
| pAG1520 | kan | mammalian | CMV-NFAST2-LifeAct-IRES2-MT2 |
| pAG1522 | kan | mammalian | CMV-LifeAct-FKBP-CFAST2-IRES2-iRFP670 |
| pAG1530 | kan | mammalian | CMV-NFAST2-MAP4-IRES2-MT2 |
| pAG1531 | kan | mammalian | CMV-MAP4-CFAST2-IRES2-iRFP670 |
| pAG1754 | kan | mammalian | CMV-CFAST2-βActin-IRES2-iRFP670 |
| pAG1936 | kan | mammalian | CMV-NFAST2-LaminAC-IRES2-MT2 |
| pAG1939 | kan | mammalian | CMV-NFAST2-BH3-IRES2-MT2 |
| pAG1967 | kan | mammalian | CMV-TOM20-Bcl-xL (ΔTMD)-CFAST2 |
| pAG1976 | kan | mammalian | CFAST2-LaminA/C-IRES2-iRFP670 |

**Supplementary Table 2. Primers**

| Number | Sequence 5' to 3' | Name |
|---|---|---|
| ag2088 | ATGATTGAACAAGATGGATTGCACGCAG | Kan-Fw |
| ag2089 | CATCTTGTTCAATCATGCGAAACGATCCTC | Kan-Rv |
| ag2194 | ctaccggactcagatctaccATGGTGTCCCGGCAAGAAG | MAP4-Fw-Gibson |
| ag2195 | ATCTGAGTCCGGTAGCGCTAGC | CMV-Rv-Gibson |
| ag2196 | tggtggcggcagctcttcgggcggagggGAGATGTGGCATGAAGGCC | FRB-Fw-Gibson |
| ag2197 | agagctgccgccaccACTAGTGTCTGGTTTAATCACACTCATGGT G | MAP4-Rv-Gibson |
| ag2202 | TTTTAGATCTACCATGGTGGGTCGGAAC | BglII-Tom20-F |
| ag2203 | AAAAACTAGTGAAGTTGGGGTCACTTCGTCTTTTG | Tom20-SpeI-R |
| ag3841 | tataagatctaccATGCCCGAACCTGCG | BglII-H2B-Fw |
| ag3842 | tataactagtCTTGGAGCTGGTGTACTTGGTC | SpeI-H2B-Rev |
| ag1858 | CAATGGGAGTTTGTTTTGGCACC | CMV-Fw |
| ag1935 | agatccgccaccctagctagcaccATGGGTTTCCCAGCCGCC | NUP133-CFAST2-Gibson |
| ag1937 | ggtggcggatctgagtccggtagcacTAGCGGATCTGACGGTTCACTAA AC | NUP133-CFAST2-Gibson |
| ag1938 | GGCTGCCTaCTAGCCTGCGACGCGC | NUP133-CFAST2-Gibson |
| ag1940 | gtAGGCAGCCGCACGGCT | iRFP670-NheI-mutagenesis |
| ag1948 | gcggcagcggcggagggtccggaATGGACAGGAGTGGATTTGGCGA | NFAST2FAST-NUP107-Gibson |
| ag1950 | agcagagcttgcagagccaccggttcaCAGCTGGATCTCATAGCCCAG AG | NFAST2FAST-NUP107-Gibson |
| ag1952 | agcagcagagcttgcagagccaccggtTCACAGTCTCTTCACGAAGAT CCAGAAGC | IRES2-Mutagenesis |
| ag1953 | aagctctgctgctggaggtATCGCCCCTCTCCCTCCC | IRES2-Mutagenesis |
| ag2088 | ATGATTGAACAAGATGGATTGCACGCAG | Kan-Fw |

| | | |
|---|---|---|
| ag2089 | CATCTTGTTCAATCATGCGAAACGATCCTC | Kan-Rv |
| ag2366 | gcagagctggtttagtgaacc | CMV promoter reverse |
| ag2367 | cggttcactaaaccagctctgctta | IRES2-Mutagenesis-2 |
| ag2368 | cctcgagcGCCCCTCTCCCTCCC | IRES2-Mutagenesis-2 |
| ag2370 | gaggggcgctcgagGATACCTCCAGCAGCAGAGCTTG | IRES2-Mutagenesis-2 |
| ag2516 | GGATTCGCtGATCCACCGGTCG | MAP4-BamHI mod-Fw |
| ag2517 | CGGTGGATCaGCGAATCCGCC | MAP4-BamHI mod-Rv |
| ag2518 | GATCCATGGGCGTGGCCGACTTGATCAAGAAGTTCGAGTC CATCTCCAAGGAGGAGTGAA | BamHI-lifeAct-AgeI-Fw |
| ag2519 | CCGGTTCACTCCTCCTTGGAGATGGACTCGAACTTCTTGAT CAAGTCGGCCACGCCCATG | BamHI-lifeAct-AgeI-Rv |
| ag2522 | GATCTGCCACCATGGGCGACAGCTTCTGGATCTTCGTGAA GAGACTGT | BglII-CFAST2-BspEI-Fw |
| ag2523 | CCGGACAGTCTCTTCACGAAGATCCAGAAGCTGTCGCCCAT GGTGGCA | BglII-CFAST2-BspEI-Rv |
| ag2557 | tatagctagcaccATGGTGTCCCGGCAAGAAG | NheI-Map4-Fw |
| ag2558 | tataggatccGTCTGGTTTAATCACACTCATGGTGG | MAP4-BamHI-Rv |
| ag2560 | tataaccggttcattaGTCTGGTTTAATCACACTCATGGTGG | MAP4-Stop-AgeI-Rv |
| ag2561 | GGTGCCAAAACAAACTCCCATTG | cmv-5'-rev |
| ag2562. | TTTCCACCTGCACTCCCCCTCCGCCCGAAG | LifeAct-linker-FKBP-rev |
| ag2563 | ggGGAGTGCAGGTGGAAACCATCTC | FKBP-Fw |
| ag2674 | TGATCCACCcGTCGCCACCATG | MAP4-AgeImut-Fw |
| ag2675 | GACGGgTGGATCAGCGAATCCG | MAP4-AgeImut-Rv |
| ag2676 | tataggatccATGGTGTCCCGGCAAGAAG | MAP4-BamHI-Fw |
| ag3723. | GCTGTTCCGACCCACCAT | TOM20 VAMPB Gibson |
| ag3732 | gagggGGATCCGAGACCCCGTC | LaminA/C AgeI mutagenesis |

| | | |
|---|---|---|
| ag3733 | CTTTGGTGGGAAgCGGTAAGTCAGC | LaminA/C AgeI mutagenesis |
| ag3734 | GCTGACTTACCGcTTCCCACCAAAG | LaminA/C AgeI mutagenesis |
| ag3735 | CGTGCAATCCATCTTGTTCAATC | Kan |
| ag3736 | GATTGAACAAGATGGATTGCACG | Kan |
| ag3737 | gacggggtctcGGATCCCCCTCCGC | NFAST2 |
| ag3758 | GATCCACCATGGGGCAGGTGGGACGGCAGCTCGCCATCAT CGGGGACGACATCAACCGATAATGAA | BH3 domain annealing AgeI |
| ag3759 | CCGGTTCATTATCGGTTGATGTCGTCCCCGATGATGGCGAG CTGCCGTCCCACCTGCCCCATGGTG | BH3 domain annealing AgeI |
| ag3762 | ATGGTGGGTCGGAACAGC | Tom20 Bcl xL Gibson |
| ag3764 | gtggcggcagctctATGAGCCAAAGCAACCG | Tom20 Bcl xL Gibson |
| ag3765 | ctttggctcatAGAGCTGCCGCCACC | Tom20 Bcl xL Gibson |
| ag3766 | gaaagggccaggaacgcGGATCCGGTGGCG | Tom20 Bcl xL Gibson |
| ag3767 | ggatccGCGTTCCTGGCCCTTTC | Tom20 Bcl xL Gibson |
| ag3818 | cagaactgcagcatcatgTAATGAACCGGTGGCTCTG | LaminA/C AgeI mutagenesis |
| ag3819 | cagagccaccggtTCATTACATGATGCTGCAGTTCTG | LaminA/C AgeI mutagenesis |

### Supplementary Table 3. Sequences

| ORF | Nucleotide sequence | Translated sequence |
|---|---|---|
| NFAST2 | atggagaccgtgagattcggcggcgacgacatcgagaacagcctggccaa<br>gatggacgacaaggccctggacaagctggccttcggcgccatccagctgga<br>cggcaacggcaagatcatccactacaacgccgccgagggcaccatcaccg<br>gcagagaccccaagaccgtgatcggcaagaacttcttcaccgacgtggccc<br>ccggcacccagagcaaggagttccagggcagattcaaggagggcgtgcag<br>aagggcgacctgaacaccatgttcgagtggatgatccccaccagcagaggc<br>cccaccaaggtgaaggtgcacatgaagaaggccatgacc | METVRFGGDDIENSLAKMD<br>DKALDKLAFGAIQLDGNGKI<br>IHYNAAEGTITGRDPKTVIG<br>KNFFTDVAPGTQSKEFQGR<br>FKEGVQKGDLNTMFEWMIP<br>TSRGPTKVKVHMKKAMT |
| CFAST2 | Ggcgacagcttctggatcttcgtgaagagactg | GDSFWIFVKRL |
| FRB | gagatgtggcatgaaggcctggaagaggcatctcgtttgtactttggggaaag<br>gaacgttaaaggcatgtttgaggtgctggagcccttgcatgctatgatggaacg<br>gggcccccagactctgaaggaaacatcctttaatcaggcctatggtcgagattt<br>aatggaggcccaagagtggtgcaggaagtacatgaaatcagggaatgtcaa<br>ggacctcacccaagcctgggacctctattatcatgtgttccgacgaatctcaaa<br>gcaggtc | EMWHEGLEEASRLYFGER<br>NVKGMFEVLEPLHAMMER<br>GPQTLKETSFNQAYGRDLM<br>EAQEWCRKYMKSGNVKDL<br>TQAWDLYYHVFRRISKQV |
| FKBP | ggagtgcaggtggaaaccatctccccaggagacgggcgcaccttccccaag<br>cgcggccagacctgcgtggtgcactacaccgggatgcttgaagatggaaag<br>aaatttgattcctcccgggacagaaacaagccctttaagtttatgctaggcaag<br>caggaggtgatccgaggctgggaagaaggggttgcccagatgagtgtgggt<br>cagagagccaaactgactatatctccagattatgcctatggtgccactgggcac<br>ccaggcatcatcccaccacatgccactctcgtcttcgatgtggagcttctaaaac<br>tggaagaa | GVQVETISPGDGRTFPKRG<br>QTCVVHYTGMLEDGKKFD<br>SSRDRNKPFKFMLGKQEVI<br>RGWEEGVAQMSVGQRAKL<br>TISPDYAYGATGHPGIIPPHA<br>TLVFDVELLKLEE |
| MAP4 | atggtgtcccggcaagaagaagcaaaggctgctgtaggtgtgactggaaatg<br>acatcactaccccgccaaacaaggagccaccaccaagcccagaaaagaa<br>agcaaagcctttggccaccactcaacctgcaaagacttcaacatcgaaagcc<br>aaaacacagcccacttctctccctaagcaaccagctcccaccacctctggtgg<br>gttgaataaaaaacccatgagcctcgcctcaggctcagtgccagctgcccca<br>cacaaacgccctgctgctgccactgctactgccaggccttccaccctacctgcc<br>agagacgtgaagccaaagccaattacagaagctaaggttgccgaaaagcg<br>gacctctccatccaagccttcatctgccccagccctcaaacctggacctaaa<br>ccaccccaaccgtttcaaaagccacatctccctcaactcttgtttccactggacc<br>aagtagtagaagtccagctacaactctgcctaagaggccaaccagcatcaa<br>gactgaggggaaacctgctgatgtcaaaaggatgactgctaagtctgcctcag<br>ctgacttgagtcgctcaaagaccacctctgccagttctgtgaagagaaacacc<br>actcccactggggcagcaccccccagcagggatgacttccactcgagtcaagc<br>ccatgtctgcacctagccgctcttctggggctctttctgtggacaagaagcccac<br>ttccactaagcctagctcctctgctcccagggtgagccgcctggccacaactgtt<br>tctgcccctgacctgaagagtgttcgctccaaggtcggctctacagaaaacatc<br>aaacaccagcctggaggaggccgggccaaggtagagaaaaaaacagag<br>gcagctaccacagctgggaagcctgaacctaatgcagtcactaaagcagcc<br>ggctccattgcgagtgcacagaaaccgcctgctgggaaagtccagatagtat<br>ccaaaaaagtgagctacagtcatattcaatccaagtgtgtttccaaggacaata<br>ttaagcatgtccctggatgtggcaatgttcagattcagaacaagaaagtggac<br>atatccaaggtctcctccaagtgtgggtccaaagctaatatcaagcacaagcct<br>ggtggaggagatgtcaagattgaaagtcagaagttgaacttcaaggagaag<br>gcccaagccaaagtgggaggcggattcgcggatccaccggtcgccaccatg<br>agtgtgattaaaccagac | MVSRQEEAKAAVGVTGNDI<br>TTPPNKEPPPSPEKKAKPL<br>ATTQPAKTSTSKAKTQPTSL<br>PKQPAPTTSGGLNKKPMSL<br>ASGSVPAAPHKRPAAATATA<br>RPSTLPARDVKPKPITEAKV<br>AEKRTSPSKPSSAPALKPG<br>PKTTPTVSKATSPSTLVSTG<br>PSSRSPATTLPKRPTSIKTE<br>GKPADVKRMTAKSASADLS<br>RSKTTSASSVKRNTTPTGA<br>APPAGMTSTRVKPMSAPSR<br>SSGALSVDKKPTSTKPSSS<br>APRVSRLATTVSAPDLKSV<br>RSKVGSTENIKHQPGGGRA<br>KVEKKTEAATTAGKPEPNA<br>VTKAAGSIASAQKPPAGKV<br>QIVSKKVSYSHIQSKCVSKD<br>NIKHVPGCGNVQIQNKKVDI<br>SKVSSKCGSKANIKHKPGG<br>GDVKIESQKLNFKEKAQAK<br>VGGGFADPPVATMSVIKPD |
| Lifeact | atgggcgtggccgacttgatcaagaagttcgagtccatctccaaggaggag | GVADLIKKFESISKEE |
| TOM20<br>(1-34) | atggtgggtcggaacagcgccatcgccgcgggcgtgtgcggtgccctcttcat<br>agggtactgcatctactttgaccgcaaaagacgaagtgaccccaacttc | MVGRNSAIAAGVCGALFIG<br>YCIYFDRKRRSDPNF |
| H2B | atgcccgaacctgcgaagtcagcgcccgctcccaaaaaaggctctaaaaaa<br>gctgtcgccaagacccagaagaaggggggataagaaaaggcgtaagacca<br>ggaaagagagttacgccatttacgtgtacaaagtactaaaacaagtccaccc<br>ggacactggcatctcctcaaaggcgatgggcattatgaactcatttgtaaacga<br>catcttcgagcgcatcgccggagaagcgtcgcgcctggcgcattacaacaag<br>cgctccactatcacatcccgggagatccagacggccgtgcgcctgctcctgcc | MPEPAKSAPAPKKGSKKAV<br>AKTQKKGDKKRRKTRKESY<br>AIYVYKVLKQVHPDTGISSK<br>AMGIMNSFVNDIFERIAGEA<br>SRLAHYNKRSTITSREIQTA |

| | | |
|---|---|---|
| | cggagaactggccaaacacgctgtgtctgagggcacaaaggccgtgacca agtacaccagctccaag | VRLLLPGELAKHAVSEGTK AVTKYTSSK |
| MT2 | atggtgagcaagggcgaggagctgttcaccggggtggtgcccatcctggtcg agctggacggcgacgtaaacggccacaagttcagcgtgtccggcgagggc gagggcgatgccacctacggcaagctgaccctgaagttcatctgcaccaccg gcaagctgcccgtgccctggcccaccctcgtgaccaccctgtcctggggcgtg cagtgcttcgcccgctaccccgaccacatgaagcagcacgacttcttcaagtc cgccatgcccgaaggctacgtccaggagcgcaccatcttcttcaaggacgac ggcaactacaagacccgcgccgaggtgaagttcgagggcgacaccctggtg aaccgcatcgagctgaagggcatcgacttcaaggaggacggcaacatcctg gggcacaagctggagtacaactacttcagcgacaacgtctatatcaccgccg acaagcagaagaacggcatcaaggccaacttcaagatccgccacaacatc gaggacggcggcgtgcagctcgccgaccactaccagcagaacacccccat cggcgacggccccgtgctgctgcccgacaaccactacctgagcacccagtc caagctgagcaaagaccccaacgagaagcgcgatcacatggtcctgctgga gttcgtgaccgccgccgggatcactctcggcatggacgagctgtacaagtaa | MVSKGEELFTGVVPILVELD GDVNGHKFSVSGEGEGDA TYGKLTLKFICTTGKLPVPW PTLVTTLSWGVQCFARYPD HMKQHDFFKSAMPEGYVQ ERTIFFKDDGNYKTRAEVKF EGDTLVNRIELKGIDFKEDG NILGHKLEYNYFSDNVYITA DKQKNGIKANFKIRHNIEDG GVQLADHYQQNTPIGDGPV LLPDNHYLSTQSKLSKDPN EKRDHMVLLEFVTAAGITLG MDELYK |
| iRFP670 | Atggcgcgtaaggtcgatctcacctcctgcgatcgcgagccgatccacatccc cggcagcattcagccgtgcggctgcctgctagcctgcgacgcgcaggcggtg cggatcacgcgcattacggaaaatgccggcgcgttctttggacgcgaaactcc gcgggtcggtgagctactcgccgattacttcggcgagaccgaagcccatgcg ctgcgcaacgcactggcgcagtcttccgatccaaagcgaccggcgctgatctt cggttggcgcgacggcctgaccggccgcaccttcgacatctcactgcatcgcc atgacggtacatcgatcatcgagttcgagcctgcggcggccgaacaggccga caatccgctgcggctgacgcggcagatcatcgcgcgcaccaaagaactgaa gtcgctcgaagagatggccgcacgggtgccgcgctatctgcaggcgatgctc ggctatcaccgcgtgatgttgtaccgcttcgcggacgacggctccgggatggtg atcggcgaggcgaagcgcagcgacctcgagagctttctcggtcagcactttcc ggcgtcgctggtcccgcagcaggcgcggctactgtacttgaagaacgcgatc cgcgtggtctcggattcgcgcggcatcagcagccggatcgtgcccgagcacg acgcctccggcgccgcgctcgatctgtcgttcgcgcacctgcgcagcatctcg ccctgccatctcgaatttctgcggaacatgggcgtcagcgcctcgatgtcgctgt cgatcatcattgacggcacgctatggggattgatcatctgtcatcattacgagcc gcgtgccgtgccgatggcgcagcgcgtcgcggccgaaatgttcgccgacttct tatcgctgcacttcaccgccgcccaccaccaacgc | MARKVDLTSCDREPIHIPGS IQPCGCLLACDAQAVRITRIT ENAGAFFGRETPRVGELLA DYFGETEAHALRNALAQSS DPKRPALIFGWRDGLTGRT FDISLHRHDGTSIIEFEPAAA EQADNPLRLTRQIIARTKEL KSLEEMAARVPRYLQAMLG YHRVMLYRFADDGSGMVIG EAKRSDLESFLGQHFPASL VPQQARLLYLKNAIRVVSDS RGISSRIVPEHDASGAALDL SFAHLRSISPCHLEFLRNMG VSASMSLSIIIDGTLWGLIIC HHYEPRAVPMAQRVAAEM FADFLSLHFTAAHHQR |
| IRES2 | Cccctctccctcccccccccctaacgttactggccgaagccgcttggaataag gccggtgtgcgtttgtctatatgttattttccaccatattgccgtcttttggcaatgtga gggcccggaaacctggccctgtcttcttgacgagcattcctaggggtctttcccc tctcgccaaaggaatgcaaggtctgttgaatgtcgtgaaggaagcagttcctct ggaagcttcttgaagacaaacaacgtctgtagcgaccctttgcaggcagcgg aaccccccacctggcgacaggtgcctctgcggccaaaagccacgtgtataa gatacacctgcaaaggcggcacaaccccagtgccacgttgtgagttggatag ttgtggaaagagtcaaatggctctcctcaagcgtattcaacaaggggctgaag gatgcccagaaggtaccccattgtatgggatctgatctggggcctcggtacac atgctttacatgtgtttagtcgaggttaaaaaaacgtctaggccccccgaacca cggggacgtggttttcctttgaaaaacacgatgataatatggccacaacc | |
| BH3 domain of BAK1 | gggcaggtgggacggcagctcgccatcatcggggacgacatcaaccga | GQVGRQLAIIGDDINR |
| Bcl xL ΔTMD | Atgagccaaagcaaccgggagctggtggttgactttctctcctacaagctttcc cagaaaggatacagctggagtcagtttagtgatgtggaagagaacaggactg aggccccagaagggactgaatcggagatggagacccccagtgccatcaat ggcaacccatcctggcacctggcagacagccccgcggtgaatggagccact ggccacagcagcagtttggatgcccgggaggtgatccccatggcagcagta aagcaagcgctgagggaggcaggcgacgagtttgaactgcggtaccggcg ggcattcagtgacctgacatcccagctccacatcaccccagggacagcatatc agagctttgaacaggtagtgaatgaactcttccgggatggggtaaactggggt cgcattgtggcctttttctccttcggcggggcactgtgcgtggaaagcgtagaca aggagatgcaggtattggtgagtcggatcgcagcttggatggccacttacctga atgaccacctagagccttggattcaggagaacggcggctgggatacttttgtgg | MSQSNRELVVDFLSYKLSQ KGYSWSQFSDVEENRTEA PEGTESEMETPSAINGNPS WHLADSPAVNGATGHSSSL DAREVIPMAAVKQALREAG DEFELRYRRAFSDLTSQLHI TPGTAYQSFEQVVNELFRD GVNWGRIVAFFSFGGALCV ESVDKEMQVLVSRIAAWMA TYLNDHLEPWIQENGGWD |

| | | |
|---|---|---|
| | aactctatgggaacaatgcagcagccgagagccgaaagggccaggaacg<br>c | TFVELYGNNAAAESRKGQE<br>R |
| $\beta$-Actin | gatgatgatatcgccgcgctcgtcgtcgacaacggctccggcatgtgcaaggc<br>cggcttcgcgggcgacgatgccccccgggccgtcttcccctccatcgtggggc<br>gccccaggcaccagggcgtgatggtgggcatgggtcagaaggattcctatgt<br>gggcgacgaggcccagagcaagagaggcatcctcaccctgaagtacccca<br>tcgagcacggcatcgtcaccaactgggacgacatggagaaaatctggcacc<br>acaccttctacaatgagctgcgtgtggctcccgaggagcaccccgtgctgctg<br>accgaggccccctgaaccccaaggccaaccgcgagaagatgacccagat<br>catgtttgagaccttcaacaccccagccatgtacgttgctatccaggctgtgctat<br>ccctgtacgcctctggccgtaccactggcatcgtgatggactccggtgacggg<br>gtcacccacactgtgcccatctacgaggggtatgccctcccccatgccatcctg<br>cgtctggacctggctggccgggacctgactgactacctcatgaagatcctcac<br>cgagcgcggctacagcttcaccaccacggccgagcgggaaatcgtgcgtga<br>cattaaggagaagctgtgctacgtcgccctggacttcgagcaagagatggcc<br>acggctgcttccagctcctccctggagaagagctacgagctgcctgacggcc<br>aggtcatcaccattggcaatgagcggttccgctgccctgaggcactcttccagc<br>cttccttcctgggcatggagtcctgtggcatccacgaaactaccttcaactccatc<br>atgaagtgtgacgtggacatccgcaaagacctgtacgccaacacagtgctgt<br>ctggcggcaccaccatgtaccctggcattgccgacaggatgcagaaggagat<br>cactgccctggcacccagcacaatgaagatcaagatcattgctcctcctgagc<br>gcaagtactccgtgtggatcggcggctccatcctggcctcgctgtccaccttcca<br>gcagatgtggatcagcaagcaggagtatgacgagtccggcccctccatcgtc<br>caccgcaaatgcttc | DDDIAALVVDNGSGMCKAG<br>FAGDDAPRAVFPSIVGRPR<br>HQGVMVGMGQKDSYVGD<br>EAQSKRGILTLKYPIEHGIVT<br>NWDDMEKIWHHTFYNELR<br>VAPEEHPVLLTEAPLNPKAN<br>REKMTQIMFETFNTPAMYV<br>AIQAVLSLYASGRTTGIVMD<br>SGDGVTHTVPIYEGYALPH<br>AILRLDLAGRDLTDYLMKILT<br>ERGYSFTTTAEREIVRDIKE<br>KLCYVALDFEQEMATAASS<br>SSLEKSYELPDGQVITIGNE<br>RFRCPEALFQPSFLGMESC<br>GIHETTFNSIMKCDVDIRKD<br>LYANTVLSGGTTMYPGIADR<br>MQKEITALAPSTMKIKIIAPP<br>ERKYSVWIGGSILASLSTFQ<br>QMWISKQEYDESGPSIVHR<br>KCF |
| Lamin<br>A/C | gagaccccgtcccagcggcgcgccacccgcagcggggcgcaggccagct<br>ccactccgctgtcgcccacccgcatcacccggctgcaggagaaggaggacc<br>tgcaggagctcaatgatcgcttggcggtctacatcgaccgtgtgcgctcgctgg<br>aaacggagaacgcagggctgcgccttcgcatcaccgagtctgaagaggtgg<br>tcagccgcgaggtgtccggcatcaaggccgcctacgaggccgagctcgggg<br>atgcccgcaagacccttgactcagtagccaaggagcgcgcccgcctgcagct<br>ggagctgagcaaagtgcgtgaggagtttaaggagctgaaagcgcgcaatac<br>caagaaggagggtgacctgatagctgctcaggctcggctgaaggacctgga<br>ggctctgctgaactccaaggaggccgcactgagcactgctctcagtgagaag<br>cgcacgctggagggcgagctgcatgatctgcggggccaggtggccaagctt<br>gaggcagccctaggtgaggccaagaagcaacttcaggatgagatgctgcgg<br>cgggtggatgctgagaacaggctgcagaccatgaaggaggaactggacttc<br>cagaagaacatctacagtgaggagctgcgtgagaccaagcgccgtcatgag<br>acccgactggtggagattgacaatgggaagcagcgtgagtttgagagccggc<br>tggcggatgcgctgcaggaactgcgggcccagcatgaggaccaggtggag<br>cagtataagaaggagctggagaagacttattctgccaagctggacaatgcca<br>ggcagtctgctgagaggaacagcaacctggtgggggctgcccacgaggag<br>ctgcagcagtcgcgcatccgcatcgacagcctctctgcccagctcagccagct<br>ccagaagcagctggcagccaaggaggcgaagcttcgagacctggaggact<br>cactggcccgtgagcgggacaccagccggcggctgctggcggaaaaggag<br>cgggagatggccgagatgcgggcaaggatgcagcagcagctggacgagta<br>ccaggagcttctggacatcaagctggccctggacatggagatccacgcctac<br>cgcaagctcttggagggcgaggaggagaggctacgcctgtcccccagccct<br>acctcgcagcgcagccgtggccgtgcttcctctcactcatcccagacacaggg<br>tgggggcagcgtcaccaaaaagcgcaaactggagtccactgagagccgca<br>gcagcttctcacagcacgcacgcactagcgggcgcgtggccgtggaggagg<br>tggatgaggagggcaagtttgtccggctgcgcaacaagtccaatgaggacca<br>gtccatgggcaattggcagatcaagcgccagaatggagatgatcccttgctga<br>cttaccgcttcccaccaaagttcaccctgaaggctgggcaggtggtgacgatct<br>gggctgcaggagctggggccacccacagccccccctaccgacctggtgtgga<br>aggcacagaacacctggggctgcgggaacagcctgcgtacggctctcatca<br>actccactggggaagaagtggccatgcgcaagctggtgcgctcagtgactgt<br>ggttgaggacgacgaggatgaggatggagatgacctgctccatcaccacca<br>cggctcccactgcagcagctcgggggaccccgctgagtacaacctgcgctcg<br>cgcaccgtgctgtgcgggacctgcgggcagcctgccgacaaggcatctgcc | ETPSQRRATRSGAQASSTP<br>LSPTRITRLQEKEDLQELND<br>RLAVYIDRVRSLETENAGLR<br>LRITESEEVVSREVSGIKAA<br>YEAELGDARKTLDSVAKER<br>ARLQLELSKVREEFKELKA<br>RNTKKEGDLIAAQARLKDLE<br>ALLNSKEAALSTALSEKRTL<br>EGELHDLRGQVAKLEAALG<br>EAKKQLQDEMLRRVDAEN<br>RLQTMKEELDFQKNIYSEEL<br>RETKRRHETRLVEIDNGKQ<br>REFESRLADALQELRAQHE<br>DQVEQYKKELEKTYSAKLD<br>NARQSAERNSNLVGAAHEE<br>LQQSRIRIDSLSAQLSQLQK<br>QLAAKEAKLRDLEDSLARE<br>RDTSRRLLAEKEREMAEMR<br>ARMQQQLDEYQELLDIKLA<br>LDMEIHAYRKLLEGEEERLR<br>LSPSPTSQRSRGRASSHSS<br>QTQGGGSVTKKRKLESTES<br>RSSFSQHARTSGRVAVEEV<br>DEEGKFVRLRNKSNEDQS<br>MGNWQIKRQNGDDPLLTY<br>RFPPKFTLKAGQVVTIWAA<br>GAGATHSPPTDLVWKAQNT<br>WGCGNSLRTALINSTGEEV<br>AMRKLVRSVTVVEDDEDED<br>GDDLLHHHHGSHCSSSGD<br>PAEYNLRSRTVLCGTCGQP<br>ADKASASGSGAQVGGPISS<br>GSSASSVTVTRSYRSVGGS<br>GGGSFGDNLVTRSYLLGNS<br>SPRTQSPQNCSIM |

| | | |
|---|---|---|
| | agcggctcaggagcccaggtgggcggacccatctcctctggctcttctgcctcc agtgtcacggtcactcgcagctaccgcagtgtggggggcagtgggggtggca gcttcggggacaatctggtcacccgctcctacctcctgggcaactccagcccc cgaacccagagcccccagaactgcagcatcatg | |
| iRFP670 (NheI mutant) | Atggcgcgtaaggtcgatctcacctcctgcgatcgcgagccgatccacatccc cggcagcattcagccgtgcggctgcctactagcctgcgacgcgcaggcggtg cggatcacgcgcattacggaaaatgccggcgcgttctttggacgcgaaactcc gcgggtcggtgagctactcgccgattacttcggcgagaccgaagcccatgcg ctgcgcaacgcactggcgcagtcctccgatccaaagcgaccggcgctgatctt cggttggcgcgacggcctgaccggccgcaccttcgacatctcactgcatcgcc atgacggtacatcgatcatcgagttcgagcctgcggcggccgaacaggccga caatccgctgcggctgacgcggcagatcatcgcgcgcaccaaagaactgaa gtcgctcgaagagatggccgcacgggtgccgcgctatctgcaggcgatgctc ggctatcaccgcgtgatgttgtaccgcttcgcggacgacggctccgggatggtg atcggcgaggcgaagcgcagcgacctcgagagctttctcggtcagcactttcc ggcgtcgctggtcccgcagcaggcgcggctactgtacttgaagaacgcgatc cgcgtggtctcggattcgcgcggcatcagcagccggatcgtgcccgagcacg acgcctccggcgccgcgctcgatctgtcgttcgcgcacctgcgcagcatctcg ccctgccatctcgaatttctgcggaacatgggcgtcagcgcctcgatgtcgctgt cgatcatcattgacggcacgctatggggattgatcatctgtcatcattacgagcc gcgtgccgtgccgatggcgcagcgcgtcgcggccgaaatgttcgccgacttct tatcgctgcacttcaccgccgcccaccaccaacgc | MARKVDLTSCDREPIHIPGS IQPCGCLLACDAQAVRITRIT ENAGAFFGRETPRVGELLA DYFGETEAHALRNALAQSS DPKRPALIFGWRDGLTGRT FDISLHRHDGTSIIEFEPAAA EQADNPLRLTRQIIARTKEL KSLEEMAARVPRYLQAMLG YHRVMLYRFADDGSGMVIG EAKRSDLESFLGQHFPASL VPQQARLLYLKNAIRVVSDS RGISSRIVPEHDASGAALDL SFAHLRSISPCHLEFLRNMG VSASMSLSIIIDGTLWGLIIC HHYEPRAVPMAQRVAAEM FADFLSLHFTAAHHQR |

### Supplementary Table 4. Open reading frames

| Plasmid | Description | ORF sequence |
|---|---|---|
| pAG1356 | MAP4-FRB-NFAST2-*IRES2*-HA-MT2 | **atg**gtgtcccggcaagaagaagcaaaggctgctgtaggtgtgactggaaatgacat<br>cactaccccgccaaacaaggagccaccaccaagcccagaaaagaaagcaaag<br>cctttggccaccactcaacctgcaaagacttcaacatcgaaagccaaaacacagc<br>ccacttctctccctaagcaaccagctcccaccacctctggtgggttgaataaaaaacc<br>catgagcctcgcctcaggctcagtgccagctgccccacacaaacgccctgctgctg<br>ccactgctactgccaggccttccaccctacctgccagagacgtgaagccaaagcca<br>attacagaagctaaggttgccgaaaagcggacctctccatccaagccttcatctgcc<br>ccagccctcaaacctggacctaaaaccaccccaaccgtttcaaaagccacatctcc<br>ctcaactcttgtttccactggaccaagtagtagaagtccagctacaactctgcctaaga<br>ggccaaccagcatcaagactgaggggaaacctgctgatgtcaaaaggatgactgc<br>taagtctgcctcagctgacttgagtcgctcaaagaccacctctgccagttctgtgaaga<br>gaaacaccactcccactggggcagcacccccagcagggatgacttccactcgagt<br>caagcccatgtctgcacctagccgctcttctggggctctttctgtggacaagaagccc<br>acttccactaagcctagctcctctgctcccagggtgagccgcctggccacaactgtttc<br>tgcccctgacctgaagagtgttcgctccaaggtcggctctacagaaaacatcaaaca<br>ccagcctggaggaggccgggccaaggtagagaaaaaaacagaggcagctacc<br>acagctgggaagcctgaacctaatgcagtcactaaagcagccggctccattgcga<br>gtgcacagaaaccgcctgctgggaaagtccagatagtatccaaaaaagtgagcta<br>cagtcatattcaatccaagtgtgtttccaaggacaatattaagcatgtccctggatgtgg<br>caatgttcagattcagaacaagaaagtggacatatccaaggtctcctccaagtgtgg<br>gtccaaagctaatatcaagcacaagcctggtggaggagatgtcaagattgaaagtc<br>agaagttgaacttcaaggagaaggcccaagccaaagtgggaggcggattcgcgg<br>atccaccggtcgccaccatgagtgtgattaaaccagacactagtggtggcggcagc<br>tcttcgggcggaggggagatgtggcatgaaggcctggaagaggcatctcgtttgtac<br>tttggggaaaggaacgttaaaggcatgtttgaggtgctggagcccttgcatgctatgat<br>ggaacggggcccccagactctgaaggaaacatcctttaatcaggcctatggtcgag<br>atttaatggaggcccaagagtggtgcaggaagtacatgaaatcagggaatgtcaag<br>gacctcacccaagcctgggacctctattatcatgtgttccgacgaatctcaaagcagg<br>tctccggaggaggcggcagcggcggaggggatccatggagaccgtgagattcg<br>gcggcgacgacatcgagaacagcctggccaagatggacgacaaggccctggac<br>aagctggccttcggcgccatccagctggacggcaacggcaagatcatccactaca<br>acgccgccgagggcaccatcaccggcagagaccccaagaccgtgatcggcaag<br>aacttcttcaccgacgtggcccccggcacccagagcaaggagttccagggcagatt<br>caaggagggcgtgcagaagggcgacctgaacaccatgttcgagtggatgatcccc<br>accagcagaggccccaccaaggtgaaggtgcacatgaagaaggccatgaccta<br>aatggccacaaccatgcgatcgtacccatacgatgttccagattacgctgaattc**atg**<br>gtgagcaagggcgaggagctgttcaccggggtggtgcccatcctggtcgagctgga<br>cggcgacgtaaacggccacaagttcagcgtgtccggcgagggcgagggcgatgc<br>cacctacggcaagctgaccctgaagttcatctgcaccaccggcaagctgcccgtgc<br>cctggcccaccctcgtgaccaccctgtcctggggcgtgcagtgcttcgcccgctaccc<br>cgaccacatgaagcagcacgacttcttcaagtccgccatgcccgaaggctacgtcc<br>aggagcgcaccatcttcttcaaggacgacggcaactacaagacccgcgccgaggt<br>gaagttcgagggcgacaccctggtgaaccgcatcgagctgaagggcatcgacttc<br>aaggaggacggcaacatcctggggcacaagctggagtacaactacttcagcgac<br>aacgtctatatcaccgccgacaagcagaagaacggcatcaaggccaacttcaag<br>atccgccacaacatcgaggacggcggcgtgcagctcgccgaccactaccagcag<br>aacacccccatcggcgacggccccgtgctgctgcccgacaaccactacctgagca<br>cccagtccaagctgagcaaagaccccaacgagaagcgcgatcacatggtcctgct<br>ggagttcgtgaccgccgccgggatcactctcggcatggacgagctgtacaagtaa |

| | | |
|---|---|---|
| pAG1359 | TOM20-FRB-NFAST2-*IRES2*-HA-MT2 | **atg**gtgggtcggaacagcgccatcgccgcgggcgtgtgcggtgccctcttcatagg<br>gtactgcatctactttgaccgcaaaagacgaagtgaccccaacttcactagtggtgg<br>cggcagctcttcgggcggagggggagatgtggcatgaaggcctggaagaggcatct<br>cgtttgtactttggggaaaggaacgttaaaggcatgtttgaggtgctggagcccttgca<br>tgctatgatggaacggggcccccagactctgaaggaaacatcctttaatcaggccta<br>tggtcgagatttaatggaggcccaagagtggtgcaggaagtacatgaaatcaggga<br>atgtcaaggacctcacccaagcctgggacctctattatcatgtgttccgacgaatctca<br>aagcaggtctccggaggaggcggcagcggcggagggggatccatggagaccgt<br>gagattcggcggcgacgacatcgagaacagcctggccaagatggacgacaagg<br>ccctggacaagctggccttcggcgccatccagctggacggcaacggcaagatcat<br>ccactacaacgccgccgagggcaccatcaccggcagagaccccaagaccgtgat<br>cggcaagaacttcttcaccgacgtggcccccggcacccagagcaaggagttccag<br>ggcagattcaaggagggcgtgcagaagggcgacctgaacaccatgttcgagtgg<br>atgatccccaccagcagaggccccaccaaggtgaaggtgcacatgaagaaggcc<br>atgacctaaatggccacaaccatgcgatcgtacccatacgatgttccagattacgctg<br>aattc**atg**gtgagcaagggcgaggagctgttcaccggggtggtgcccatcctggtc<br>gagctggacggcgacgtaaacggccacaagttcagcgtgtccggcgagggcgag<br>ggcgatgccacctacggcaagctgaccctgaagttcatctgcaccaccggcaagct<br>gcccgtgccctggcccaccctcgtgaccaccctgtcctggggcgtgcagtgcttcgc<br>ccgctaccccgaccacatgaagcagcacgacttcttcaagtccgccatgcccgaag<br>gctacgtccaggagcgcaccatcttcttcaaggacgacggcaactacaagacccg<br>cgccgaggtgaagttcgagggcgacaccctggtgaaccgcatcgagctgaaggg<br>catcgacttcaaggaggacggcaacatcctggggcacaagctggagtacaactac<br>ttcagcgacaacgtctatatcaccgccgacaagcagaagaacggcatcaaggcca<br>acttcaagatccgccacaacatcgaggacggcggcgtgcagctcgccgaccacta<br>ccagcagaacacccccatcggcgacggccccgtgctgctgcccgacaaccacta<br>cctgagcacccagtccaagctgagcaaagaccccaacgagaagcgcgatcacat<br>ggtcctgctggagttcgtgaccgccgccgggatcactctcggcatggacgagctgta<br>caagtaa |
| pAG1965 | H2B-FRB-NFAST2-*IRES2*-HA-MT2 | **atg**cccgaacctgcgaagtcagcgcccgctcccaaaaaaggctctaaaaaagct<br>gtcgccaagacccagaagaaggggggataagaaaaggcgtaagaccaggaaag<br>agagttacgccatttacgtgtacaaagtactaaaacaagtccacccggacactggc<br>atctcctcaaaggcgatgggcattatgaactcatttgtaaacgacatcttcgagcgcat<br>cgccggagaagcgtcgcgcctggcgcattacaacaagcgctccactatcacatccc<br>gggagatccagacggccgtgcgcctgctcctgcccggagaactggccaaacacg<br>ctgtgtctgagggcacaaaggccgtgaccaagtacaccagctccaagactagtggt<br>ggcggcagctcttcgggcggagggggagatgtggcatgaaggcctggaagaggca<br>tctcgtttgtactttggggaaaggaacgttaaaggcatgtttgaggtgctggagcccttg<br>catgctatgatggaacggggcccccagactctgaaggaaacatcctttaatcaggc<br>ctatggtcgagatttaatggaggcccaagagtggtgcaggaagtacatgaaatcag<br>ggaatgtcaaggacctcacccaagcctgggacctctattatcatgtgttccgacgaat<br>ctcaaagcaggtctccggaggaggcggcagcggcggagggggatccatggaga<br>ccgtgagattcggcggcgacgacatcgagaacagcctggccaagatggacgaca<br>aggccctggacaagctggccttcggcgccatccagctggacggcaacggcaagat<br>catccactacaacgccgccgagggcaccatcaccggcagagaccccaagaccgt<br>gatcggcaagaacttcttcaccgacgtggcccccggcacccagagcaaggagttc<br>cagggcagattcaaggagggcgtgcagaagggcgacctgaacaccatgttcgagt<br>ggatgatccccaccagcagaggccccaccaaggtgaaggtgcacatgaagaag<br>gccatgacctaaatggccacaaccatgcgatcgtacccatacgatgttccagattac<br>gctgaattc**atg**gtgagcaagggcgaggagctgttcaccggggtggtgcccatcct<br>ggtcgagctggacggcgacgtaaacggccacaagttcagcgtgtccggcgaggg<br>cgagggcgatgccacctacggcaagctgaccctgaagttcatctgcaccaccggc<br>aagctgcccgtgccctggcccaccctcgtgaccaccctgtcctggggcgtgcagtgc<br>ttcgcccgctaccccgaccacatgaagcagcacgacttcttcaagtccgccatgccc<br>gaaggctacgtccaggagcgcaccatcttcttcaaggacgacggcaactacaaga<br>cccgcgccgaggtgaagttcgagggcgacaccctggtgaaccgcatcgagctgaa<br>gggcatcgacttcaaggaggacggcaacatcctggggcacaagctggagtacaa<br>ctacttcagcgacaacgtctatatcaccgccgacaagcagaagaacggcatcaag<br>gccaacttcaagatccgccacaacatcgaggacggcggcgtgcagctcgccgac<br>cactaccagcagaacacccccatcggcgacggccccgtgctgctgcccgacaac |

| | | |
|---|---|---|
| | | cactacctgagcacccagtccaagctgagcaaagaccccaacgagaagcgcgat<br>cacatggtcctgctggagttcgtgaccgccgccgggatcactctcggcatggacgag<br>ctgtacaagtaa |
| pAG580 | C-Myc-FKBP-CFAST2-*IRES2*-HA -iRFP670-C-Myc | **atg**gaacaaaagcttatttctgaagaggacttggaattcggagtgcaggtggaaac<br>catctccccaggagacgggcgcaccttccccaagcgcggccagacctgcgtggtg<br>cactacaccgggatgcttgaagatggaaagaaatttgattcctcccgggacagaaa<br>caagccctttaagtttatgctaggcaagcaggaggtgatccgaggctgggaagaag<br>gggttgcccagatgagtgtgggtcagagagccaaactgactatatctccagattatgc<br>ctatggtgccactgggcacccaggcatcatcccaccacatgccactctcgtcttcgat<br>gtggagcttctaaaactggaagaatccggaggaggcggcagcggcggaggggg<br>atccggcgacagcttctggatcttcgtgaagagactgtaa**atg**gccacaaccatgcg<br>atcgtacccatacgatgttccagattacgctgaattc**atg**gcgcgtaaggtcgatctca<br>cctcctgcgatcgcgagccgatccacatccccggcagcattcagccgtgcggctgc<br>ctgctagcctgcgacgcgcaggcggtgcggatcacgcgcattacggaaaatgccg<br>gcgcgttctttggacgcgaaactccgcgggtcggtgagctactcgccgattacttcgg<br>cgagaccgaagcccatgcgctgcgcaacgcactggcgcagtcttccgatccaaag<br>cgaccggcgctgatcttcggttggcgcgacggcctgaccggccgcaccttcgacatc<br>tcactgcatcgccatgacggtacatcgatcatcgagttcgagcctgcggcggccgaa<br>caggccgacaatccgctgcggctgacgcggcagatcatcgcgcgcaccaaagaa<br>ctgaagtcgctcgaagagatggccgcacgggtgccgcgctatctgcaggcgatgct<br>cggctatcaccgcgtgatgttgtaccgcttcgcggacgacggctccgggatggtgatc<br>ggcgaggcgaagcgcagcgacctcgagagctttctcggtcagcactttccggcgtc<br>gctggtcccgcagcaggcgcggctactgtacttgaagaacgcgatccgcgtggtctc<br>ggattcgcgcggcatcagcagccggatcgtgcccgagcacgacgcctccggcgcc<br>gcgctcgatctgtcgttcgcgcacctgcgcagcatctcgccctgccatctcgaatttctg<br>cggaacatgggcgtcagcgcctcgatgtcgctgtcgatcatcattgacggcacgctat<br>ggggattgatcatctgtcatcattacgagccgcgtgccgtgccgatggcgcagcgcg<br>tcgcggccgaaatgttcgccgacttcttatcgctgcacttcaccgccgcccaccacca<br>acgcgaacaaaagcttatttctgaagaggacttgtaa |
| pAG657 | CMV-H2B-pFAST-C-Myc | atgcccgaacctgcgaagtcagcgcccgctcccaaaaaaggctctaaaaagctg<br>tcgccaagacccagaagaaggggataagaaaaggcgtaagaccaggaaaga<br>gagttacgccatttacgtgtacaaagtactaaaacaagtccacccggacactggcat<br>ctcctcaaaggcgatgggcattatgaactcatttgtaaacgacatcttcgagcgcatc<br>gccggagaagcgtcgcgcctggcgcattacaacaagcgctccactatcacatccc<br>gggagatccagacggccgtgcgcctgctcctgcccggagaactggccaaacacg<br>ctgtgtctgagggcacaaaggccgtgaccaagtacaccagctccaagggcggag<br>gctccggaggcggatctgccaccatggagcatgttgcctttggcagtgaggacatcg<br>agaacactctggccaatatggacgacgaacaactggataggttggcctttggcgtaa<br>ttcagctcgatggtgacgggaatatcctgctgtacaatgctgctgaaggggacatcac<br>tggcagagatcccaaacaggtgattgggaagaacttcttcaaggatgttgcacctgg<br>aacggatactcccgagttttacggcaaattcaaggaaggcgcagcgtcagggaatc<br>tgaacaccatgttcgaatggacgataccgacaagcaggggaccaaccaaggtca<br>aggtgcacttgaagaaagccctttccggtgacagatattgggtctttgtgaaacgggt<br>gggatccgaacaaaagcttatttctgaagaggacttgtaa |
| pAG1520 | NFAST2-LifeAct-IRES2-HA-MT2 | atggagaccgtgagattcggcggcgacgacatcgagaacagcctggccaagatg<br>gacgacaaggccctggacaagctggccttcggcgccatccagctggacggcaac<br>ggcaagatcatccactacaacgccgccgagggcaccatcaccggcagagacccc<br>aagaccgtgatcggcaagaacttcttcaccgacgtggcccccggcacccagagca<br>aggagttccagggcagattcaaggagggcgtgcagaagggcgacctgaacacca<br>tgttcgagtggatgatccccaccagcagaggccccaccaaggtgaaggtgcacat<br>gaagaaggccatgacctccggaggaggcggcagcggcggagggggatccatgg<br>gcgtggccgacttgatcaagaagttcgagtccatctccaaggaggagtgaatggcc<br>acaaccatgcgatcgtacccatacgatgttccagattacgctgaattcatggtgagca<br>agggcgaggagctgttcaccggggtggtgcccatcctggtcgagctggacggcga<br>cgtaaacggccacaagttcagcgtgtccggcgagggcgagggcgatgccacctac<br>ggcaagctgaccctgaagttcatctgcaccaccggcaagctgcccgtgccctggcc<br>caccctcgtgaccaccctgtcctggggcgtgcagtgcttcgcccgctaccccgacca<br>catgaagcagcacgacttcttcaagtccgccatgcccgaaggctacgtccaggagc<br>gcaccatcttcttcaaggacgacggcaactacaagacccgcgccgaggtgaagttc<br>gagggcgacaccctggtgaaccgcatcgagctgaagggcatcgacttcaaggag |

| | | |
|---|---|---|
| | | gacggcaacatcctggggcacaagctggagtacaactacttcagcgacaacgtct<br>atatcaccgccgacaagcagaagaacggcatcaaggccaacttcaagatccgcc<br>acaacatcgaggacggcggcgtgcagctcgccgaccactaccagcagaacacc<br>cccatcggcgacggccccgtgctgctgcccgacaaccactacctgagcacccagt<br>ccaagctgagcaaagaccccaacgagaagcgcgatcacatggtcctgctggagtt<br>cgtgaccgccgccgggatcactctcggcatggacgagctgtacaagtaa |
| pAG1522 | LifeAct-FKBP-CFAST2-IRES2- HA-iRFP670 | atgggcgtggccgacttgatcaagaagttcgagtccatctccaaggaggagggatc<br>cggtggcggcagcggcggagggggagtgcaggtggaaaccatctccccaggag<br>acgggcgcaccttccccaagcgcggccagacctgcgtggtgcactacaccgggat<br>gcttgaagatggaaagaaatttgattcctcccgggacagaaacaagccctttaagttt<br>atgctaggcaagcaggaggtgatccgaggctgggaagaaggggttgcccagatg<br>agtgtgggtcagagagccaaactgactatatctccagattatgcctatggtgccactg<br>ggcacccaggcatcatcccaccacatgccactctcgtcttcgatgtggagcttctaaa<br>actggaagaatccggagggggcggcagcggcggagggggatccggcgacagct<br>tctggatcttcgtgaagagactgtaaatggccacaaccatgcgatcgtacccatacg<br>atgttccagattacgctgaattcatggcgcgtaaggtcgatctcacctcctgcgatcgc<br>gagccgatccacatccccggcagcattcagccgtgcggctgcctgctagcctgcga<br>cgcgcaggcggtgcggatcacgcgcattacggaaaatgccggcgcgttctttggac<br>gcgaaactccgcgggtcggtgagctactcgccgattacttcggcgagaccgaagcc<br>catgcgctgcgcaacgcactggcgcagtcttccgatccaaagcgaccggcgctgat<br>cttcggttggcgcgacggcctgaccggccgcaccttcgacatctcactgcatcgccat<br>gacggtacatcgatcatcgagttcgagcctgcggcggccgaacaggccgacaatc<br>cgctgcggctgacgcggcagatcatcgcgcgcaccaaagaactgaagtcgctcga<br>agagatggccgcacgggtgccgcgctatctgcaggcgatgctcggctatcaccgcg<br>tgatgttgtaccgcttcgcggacgacggctccgggatggtgatcggcgaggcgaag<br>cgcagcgacctcgagagctttctcggtcagcactttccggcgtcgctggtcccgcagc<br>aggcgcggctactgtacttgaagaacgcgatccgcgtggtctcggattcgcgcggc<br>atcagcagccggatcgtgcccgagcacgacgcctccggcgccgcgctcgatctgtc<br>gttcgcgcacctgcgcagcatctcgccctgccatctcgaatttctgcggaacatgggc<br>gtcagcgcctcgatgtcgctgtcgatcatcattgacggcacgctatggggattgatcat<br>ctgtcatcattacgagccgcgtgccgtgccgatggcgcagcgcgtcgcggccgaaa<br>tgttcgccgacttcttatcgctgcacttcaccgccgcccaccaccaacgcgaacaaa<br>agcttatttctgaagaggacttgtaa |
| pAG1754 | CFAST2-βActin-IRES2-HA-iRFP670 | atgggcgacagcttctggatcttcgtgaagagactgtccggaggaggcggcagcgg<br>cggagggggatccatggatgatgatatcgccgcgctcgtcgtcgacaacggctccg<br>gcatgtgcaaggccggcttcgcgggcgacgatgccccccgggccgtcttcccctcc<br>atcgtggggcgccccaggcaccagggcgtgatggtgggcatgggtcagaaggatt<br>cctatgtgggcgacgaggcccagagcaagagaggcatcctcaccctgaagtaccc<br>catcgagcacggcatcgtcaccaactgggacgacatggagaaaatctggcaccac<br>accttctacaatgagctgcgtgtggctcccgaggagcaccccgtgctgctgaccgag<br>gccccccctgaaccccaaggccaaccgcgagaagatgacccagatcatgtttgaga<br>ccttcaacaccccagccatgtacgttgctatccaggctgtgctatccctgtacgcctctg<br>gccgtaccactggcatcgtgatggactccggtgacggggtcacccacactgtgccc<br>atctacgaggggtatgccctcccccatgccatcctgcgtctggacctggctggccgg<br>gacctgactgactacctcatgaagatcctcaccgagcgcggctacagcttcaccacc<br>acggccgagcgggaaatcgtgcgtgacattaaggagaagctgtgctacgtcgccct<br>ggacttcgagcaagagatggccacggctgcttccagctcctccctggagaagagct<br>acgagctgcctgacggccaggtcatcaccattggcaatgagcggttccgctgccctg<br>aggcactcttccagccttccttcctgggcatggagtcctgtggcatccacgaaactac<br>cttcaactccatcatgaagtgtgacgtggacatccgcaaagacctgtacgccaacac<br>agtgctgtctggcggcaccaccatgtaccctggcattgccgacaggatgcagaagg<br>agatcactgccctggcacccagcacaatgaagatcaagatcattgctcctcctgagc<br>gcaagtactccgtgtggatcggcggctccatcctggcctcgctgtccaccttccagca<br>gatgtggatcagcaagcaggagtatgacgagtccggcccctccatcgtccaccgca<br>aatgcttctaaatggccacaaccatgcgatcgtacccatacgatgttccagattacgct<br>gaattcatggcgcgtaaggtcgatctcacctcctgcgatcgcgagccgatccacatc<br>cccggcagcattcagccgtgcggctgcctactagcctgcgacgcgcaggcggtgc<br>ggatcacgcgcattacggaaaatgccggcgcgttctttggacgcgaaactccgcgg |

| | | |
|---|---|---|
| | | gtcggtgagctactcgccgattacttcggcgagaccgaagcccatgcgctgcgcaa<br>cgcactggcgcagtcctccgatccaaagcgaccggcgctgatcttcggttggcgcg<br>acggcctgaccggccgcaccttcgacatctcactgcatcgccatgacggtacatcga<br>tcatcgagttcgagcctgcggcggccgaacaggccgacaatccgctgcggctgac<br>gcggcagatcatcgcgcgcaccaaagaactgaagtcgctcgaagagatggccgc<br>acgggtgccgcgctatctgcaggcgatgctcggctatcaccgcgtgatgttgtaccgc<br>ttcgcggacgacggctccgggatggtgatcggcgaggcgaagcgcagcgacctc<br>gagagctttctcggtcagcactttccggcgtcgctggtcccgcagcaggcgcggcta<br>ctgtacttgaagaacgcgatccgcgtggtctcggattcgcgcggcatcagcagccgg<br>atcgtgcccgagcacgacgcctccggcgccgcgctcgatctgtcgttcgcgcacctg<br>cgcagcatctcgccctgccatctcgaatttctgcggaacatgggcgtcagcgcctcg<br>atgtcgctgtcgatcatcattgacggcacgctatggggattgatcatctgtcatcattac<br>gagccgcgtgccgtgccgatggcgcagcgcgtcgcggccgaaatgttcgccgactt<br>cttatcgctgcacttcaccgccgcccaccaccaacgctaa |
| pAG1530 | NFAST2-MAP4-<br>IRES2-HA-MT2 | atggagaccgtgagattcggcggcgacgacatcgagaacagcctggccaagatg<br>gacgacaaggccctggacaagctggccttcggcgccatccagctggacggcaac<br>ggcaagatcatccactacaacgccgccgagggcaccatcaccggcagagacccc<br>aagaccgtgatcggcaagaacttcttcaccgacgtggcccccggcacccagagca<br>aggagttccagggcagattcaaggagggcgtgcagaagggcgacctgaacacca<br>tgttcgagtggatgatccccaccagcagaggccccaccaaggtgaaggtgcacat<br>gaagaaggccatgacctccggaggaggcggcagcggcggagggggatccatgg<br>tgtcccggcaagaagaagcaaaggctgctgtaggtgtgactggaaatgacatcact<br>accccgccaaacaaggagccaccaccaagcccagaaaagaaagcaaagccttt<br>ggccaccactcaacctgcaaagacttcaacatcgaaagccaaaacacagcccac<br>ttctctccctaagcaaccagctcccaccacctctggtgggttgaataaaaaacccatg<br>agcctcgcctcaggctcagtgccagctgccccacacaaacgccctgctgctgccac<br>tgctactgccaggccttccaccctacctgccagagacgtgaagccaaagccaatta<br>cagaagctaaggttgccgaaaagcggacctctccatccaagccttcatctgcccca<br>gccctcaaacctggacctaaaaccaccccaaccgtttcaaaagccacatctccctc<br>aactcttgtttccactggaccaagtagtagaagtccagctacaactctgcctaagagg<br>ccaaccagcatcaagactgaggggaaacctgctgatgtcaaaaggatgactgcta<br>agtctgcctcagctgacttgagtcgctcaaagaccacctctgccagttctgtgaagag<br>aaacaccactcccactggggcagcacccccagcagggatgacttccactcgagtc<br>aagcccatgtctgcacctagccgctcttctggggctctttctgtggacaagaagccca<br>cttccactaagcctagctcctctgctcccagggtgagccgcctggccacaactgtttct<br>gcccctgacctgaagagtgttcgctccaaggtcggctctacagaaaacatcaaaca<br>ccagcctggaggaggccgggccaaggtagagaaaaaaacagaggcagctacc<br>acagctgggaagcctgaacctaatgcagtcactaaagcagccggctccattgcga<br>gtgcacagaaaccgcctgctgggaaagtccagatagtatccaaaaaagtgagcta<br>cagtcatattcaatccaagtgtgtttccaaggacaatattaagcatgtccctggatgtgg<br>caatgttcagattcagaacaagaaagtggacatatccaaggtctcctccaagtgtgg<br>gtccaaagctaatatcaagcacaagcctggtggaggagatgtcaagattgaaagtc<br>agaagttgaacttcaaggagaaggcccaagccaaagtgggaggcggattcgctg<br>atccacccgtcgccaccatgagtgtgattaaaccagactaaatggccacaaccatg<br>cgatcgtacccatacgatgttccagattacgctgaattcatggtgagcaagggcgag<br>gagctgttcaccggggtggtgcccatcctggtcgagctggacggcgacgtaaacgg<br>ccacaagttcagcgtgtccggcgagggcgagggcgatgccacctacggcaagctg<br>accctgaagttcatctgcaccaccggcaagctgcccgtgccctggcccaccctcgtg<br>accaccctgtcctggggcgtgcagtgcttcgcccgctaccccgaccacatgaagca<br>gcacgacttcttcaagtccgccatgcccgaaggctacgtccaggagcgcaccatctt<br>cttcaaggacgacggcaactacaagacccgcgccgaggtgaagttcgagggcga<br>caccctggtgaaccgcatcgagctgaagggcatcgacttcaaggaggacggcaa<br>catcctggggcacaagctggagtacaactacttcagcgacaacgtctatatcaccgc<br>cgacaagcagaagaacggcatcaaggccaacttcaagatccgccacaacatcga<br>ggacggcggcgtgcagctcgccgaccactaccagcagaacacccccatcggcga<br>cggccccgtgctgctgcccgacaaccactacctgagcacccagtccaagctgagc<br>aaagaccccaacgagaagcgcgatcacatggtcctgctggagttcgtgaccgccg<br>ccgggatcactctcggcatggacgagctgtacaagtaa |

| | | |
|---|---|---|
| pAG1531 | MAP4-CFAST2-IRES2-HA-iRFP670 | atggtgtcccggcaagaagaagcaaaggctgctgtaggtgtgactggaaatgacat<br>cactaccccgccaaacaaggagccaccaccaagcccagaaaagaaagcaaag<br>cctttggccaccactcaacctgcaaagacttcaacatcgaaagccaaaacacagc<br>ccacttctctccctaagcaaccagctcccaccacctctggtgggttgaataaaaaacc<br>catgagcctcgcctcaggctcagtgccagctgccccacacaaacgccctgctgctg<br>ccactgctactgccaggccttccaccctacctgccagagacgtgaagccaaagcca<br>attacagaagctaaggttgccgaaaagcggacctctccatccaagccttcatctgcc<br>ccagccctcaaacctggacctaaaaccaccccaaccgtttcaaaagccacatctcc<br>ctcaactcttgtttccactggaccaagtagtagaagtccagctacaactctgcctaaga<br>ggccaaccagcatcaagactgaggggaaacctgctgatgtcaaaaggatgactgc<br>taagtctgcctcagctgacttgagtcgctcaaagaccacctctgccagttctgtgaaga<br>gaaacaccactcccactggggcagcacccccagcagggatgacttccactcgagt<br>caagcccatgtctgcacctagccgctcttctggggctctttctgtggacaagaagccc<br>acttccactaagcctagctcctctgctcccagggtgagccgcctggccacaactgtttc<br>tgcccctgacctgaagagtgttcgctccaaggtcggctctacagaaaacatcaaaca<br>ccagcctggaggaggccgggccaaggtagagaaaaaaacagaggcagctacc<br>acagctgggaagcctgaacctaatgcagtcactaaagcagccggctccattgcga<br>gtgcacagaaaccgcctgctgggaaagtccagatagtatccaaaaaagtgagcta<br>cagtcatattcaatccaagtgtgtttccaaggacaatattaagcatgtccctggatgtgg<br>caatgttcagattcagaacaagaaagtggacatatccaaggtctcctccaagtgtgg<br>gtccaaagctaatatcaagcacaagcctggtggaggagatgtcaagattgaaagtc<br>agaagttgaacttcaaggagaaggcccaagccaaagtgggaggcggattcgctg<br>atccacccgtcgccaccatgagtgtgattaaaccagacggatccggtggcggcagc<br>ggcggagggtccggaggcgacagcttctggatcttcgtgaagagactgtgaatggc<br>cacaaccatgcgatcgtacccatacgatgttccagattacgctgaattcatggcgcgt<br>aaggtcgatctcacctcctgcgatcgcgagccgatccacatccccggcagcattcag<br>ccgtgcggctgcctactagcctgcgacgcgcaggcggtgcggatcacgcgcattac<br>ggaaaatgccggcgcgttctttggacgcgaaactccgcgggtcggtgagctactcg<br>ccgattacttcggcgagaccgaagcccatgcgctgcgcaacgcactggcgcagtc<br>ctccgatccaaagcgaccggcgctgatcttcggttggcgcgacggcctgaccggcc<br>gcaccttcgacatctcactgcatcgccatgacggtacatcgatcatcgagttcgagcc<br>tgcggcggccgaacaggccgacaatccgctgcggctgacgcggcagatcatcgc<br>gcgcaccaaagaactgaagtcgctcgaagagatggccgcacgggtgccgcgcta<br>tctgcaggcgatgctcggctatcaccgcgtgatgttgtaccgcttcgcggacgacggc<br>tccgggatggtgatcggcgaggcgaagcgcagcgacctcgagagctttctcggtca<br>gcactttccggcgtcgctggtcccgcagcaggcgcggctactgtacttgaagaacgc<br>gatccgcgtggtctcggattcgcgcggcatcagcagccggatcgtgcccgagcacg<br>acgcctccggcgccgcgctcgatctgtcgttcgcgcacctgcgcagcatctcgccct<br>gccatctcgaatttctgcggaacatgggcgtcagcgcctcgatgtcgctgtcgatcatc<br>attgacggcacgctatggggattgatcatctgtcatcattacgagccgcgtgccgtgc<br>cgatggcgcagcgcgtcgcggccgaaatgttcgccgacttcttatcgctgcacttcac<br>cgccgcccaccaccaacgctaa |
| pAG1936 | NFAST2-LaminA/C-IRES2- HA-MT2 | atggagaccgtgagattcggcggcgacgacatcgagaacagcctggccaagatg<br>gacgacaaggccctggacaagctggccttcggcgccatccagctggacggcaac<br>ggcaagatcatccactacaacgccgccgagggcaccatcaccggcagagacccc<br>aagaccgtgatcggcaagaacttcttcaccgacgtggcccccggcacccagagca<br>aggagttccagggcagattcaaggagggcgtgcagaagggcgacctgaacacca<br>tgttcgagtggatgatccccaccagcagaggccccaccaaggtgaaggtgcacat<br>gaagaaggccatgacctccggaggaggcggcagcggcggagggggatccgag<br>accccgtcccagcggcgcgccacccgcagcggggcgcaggccagctccactccg<br>ctgtcgcccacccgcatcacccggctgcaggagaaggaggacctgcaggagctc<br>aatgatcgcttggcggtctacatcgaccgtgtgcgctcgctggaaacggagaacgc<br>agggctgcgccttcgcatcaccgagtctgaagaggtggtcagccgcgaggtgtccg<br>gcatcaaggccgcctacgaggccgagctcggggatgcccgcaagacccttgactc<br>agtagccaaggagcgcgcccgcctgcagctggagctgagcaaagtgcgtgagga<br>gtttaaggagctgaaagcgcgcaataccaagaaggagggtgacctgatagctgct<br>caggctcggctgaaggacctggaggctctgctgaactccaaggaggccgcactga<br>gcactgctctcagtgagaagcgcacgctggagggcgagctgcatgatctgcgggg<br>ccaggtggccaagcttgaggcagccctaggtgaggccaagaagcaacttcaggat |

| | | |
|---|---|---|
| | | gagatgctgcggcgggtggatgctgagaacaggctgcagaccatgaaggaggaa<br>ctggacttccagaagaacatctacagtgaggagctgcgtgagaccaagcgccgtc<br>atgagacccgactggtggagattgacaatgggaagcagcgtgagtttgagagccg<br>gctggcggatgcgctgcaggaactgcgggcccagcatgaggaccaggtggagca<br>gtataagaaggagctggagaagacttattctgccaagctggacaatgccaggcagt<br>ctgctgagaggaacagcaacctggtgggggctgcccacgaggagctgcagcagt<br>cgcgcatccgcatcgacagcctctctgcccagctcagccagctccagaagcagctg<br>gcagccaaggaggcgaagcttcgagacctggaggactcactggcccgtgagcgg<br>gacaccagccggcggctgctggcggaaaaggagcgggagatggccgagatgcg<br>ggcaaggatgcagcagcagctggacgagtaccaggagcttctggacatcaagctg<br>gccctggacatggagatccacgcctaccgcaagctcttggagggcgaggaggag<br>aggctacgcctgtcccccagccctacctcgcagcgcagccgtggccgtgcttcctctc<br>actcatcccagacacagggtgggggcagcgtcaccaaaaagcgcaaactggagt<br>ccactgagagccgcagcagcttctcacagcacgcacgcactagcgggcgcgtgg<br>ccgtggaggaggtggatgaggagggcaagtttgtccggctgcgcaacaagtccaa<br>tgaggaccagtccatgggcaattggcagatcaagcgccagaatggagatgatccct<br>tgctgacttaccgcttcccaccaaagttcaccctgaaggctgggcaggtggtgacgat<br>ctgggctgcaggagctggggccacccacagccccccctaccgacctggtgtggaag<br>gcacagaacacctggggctgcgggaacagcctgcgtacggctctcatcaactcca<br>ctggggaagaagtggccatgcgcaagctggtgcgctcagtgactgtggttgaggac<br>gacgaggatgaggatggagatgacctgctccatcaccaccacggctcccactgca<br>gcagctcgggggaccccgctgagtacaacctgcgctcgcgcaccgtgctgtgcgg<br>gacctgcgggcagcctgccgacaaggcatctgccagcggctcaggagcccaggt<br>gggcggacccatctcctctggctcttctgcctccagtgtcacggtcactcgcagctacc<br>gcagtgtgggggggcagtgggggtggcagcttcggggacaatctggtcacccgctcc<br>tacctcctgggcaactccagcccccgaacccagagcccccagaactgcagcatca<br>tgtaaatggccacaaccatgcgatcgtacccatacgatgttccagattacgctgaatt<br>catggtgagcaagggcgaggagctgttcaccggggtggtgcccatcctggtcgagc<br>tggacggcgacgtaaacggccacaagttcagcgtgtccggcgagggcgagggcg<br>atgccacctacggcaagctgaccctgaagttcatctgcaccaccggcaagctgccc<br>gtgccctggcccaccctcgtgaccaccctgtcctggggcgtgcagtgcttcgcccgct<br>accccgaccacatgaagcagcacgacttcttcaagtccgccatgcccgaaggctac<br>gtccaggagcgcaccatcttcttcaaggacgacggcaactacaagacccgcgccg<br>aggtgaagttcgagggcgacaccctggtgaaccgcatcgagctgaagggcatcga<br>cttcaaggaggacggcaacatcctggggcacaagctggagtacaactacttcagc<br>gacaacgtctatatcaccgccgacaagcagaagaacggcatcaaggccaacttca<br>agatccgccacaacatcgaggacggcggcgtgcagctcgccgaccactaccagc<br>agaacacccccatcggcgacggccccgtgctgctgcccgacaaccactacctgag<br>cacccagtccaagctgagcaaagaccccaacgagaagcgcgatcacatggtcct<br>gctggagttcgtgaccgccgccgggatcactctcggcatggacgagctgtacaagt<br>aa |
| pAG1939 | NFAST2-BH3-<br>IRES2- HA-MT2 | atggagaccgtgagattcggcggcgacgacatcgagaacagcctggccaagatg<br>gacgacaaggccctggacaagctggccttcggcgccatccagctggacggcaac<br>ggcaagatcatccactacaacgccgccgagggcaccatcaccggcagagacccc<br>aagaccgtgatcggcaagaacttcttcaccgacgtggcccccggcacccagagca<br>aggagttccagggcagattcaaggagggcgtgcagaagggcgacctgaacacca<br>tgttcgagtggatgatccccaccagcagaggccccaccaaggtgaaggtgcacat<br>gaagaaggccatgacctccggaggaggcggcagcggcggagggggatccacc<br>atggggcaggtgggacggcagctcgccatcatcggggacgacatcaaccgataa<br>atggccacaaccatgcgatcgtacccatacgatgttccagattacgctgaattcatgg<br>tgagcaagggcgaggagctgttcaccggggtggtgcccatcctggtcgagctggac<br>ggcgacgtaaacggccacaagttcagcgtgtccggcgagggcgagggcgatgcc<br>acctacggcaagctgaccctgaagttcatctgcaccaccggcaagctgcccgtgcc<br>ctggcccaccctcgtgaccaccctgtcctggggcgtgcagtgcttcgcccgctacccc<br>gaccacatgaagcagcacgacttcttcaagtccgccatgcccgaaggctacgtcca<br>ggagcgcaccatcttcttcaaggacgacggcaactacaagacccgcgccgaggtg<br>aagttcgagggcgacaccctggtgaaccgcatcgagctgaagggcatcgacttca<br>aggaggacggcaacatcctggggcacaagctggagtacaactacttcagcgaca<br>acgtctatatcaccgccgacaagcagaagaacggcatcaaggccaacttcaagat |

| | | |
|---|---|---|
| | | ccgccacaacatcgaggacggcggcgtgcagctcgccgaccactaccagaga<br>acacccccatcggcgacggccccgtgctgctgcccgacaaccactacctgagcac<br>ccagtccaagctgagcaaagaccccaacgagaagcgcgatcacatggtcctgctg<br>gagttcgtgaccgccgccgggatcactctcggcatggacgagctgtacaagtaa |
| pAG1967 | TOM20-Bcl-xL (ΔTMD) -CFAST2-IRES2 - HA-iRFP670 | atggtgggtcggaacagcgccatcgccgcgggcgtgtgcggtgccctcttcatagg<br>gtactgcatctactttgaccgcaaaagacgaagtgaccccaacttcactagtggtgg<br>cggcagctctatgagccaaagcaaccgggagctggtggttgactttctctcctacaa<br>gctttcccagaaaggatacagctggagtcagtttagtgatgtggaagagaacagga<br>ctgaggccccagaagggactgaatcggagatggagacccccagtgccatcaatg<br>gcaacccatcctggcacctggcagacagccccgcggtgaatggagccactggcc<br>acagcagcagtttggatgcccgggaggtgatccccatggcagcagtaaagcaagc<br>gctgagggaggcaggcgacgagtttgaactgcggtaccggcgggcattcagtgac<br>ctgacatcccagctccacatcaccccagggacagcatatcagagctttgaacaggt<br>agtgaatgaactcttccgggatggggtaaactggggtcgcattgtggcctttttctccttc<br>ggcggggcactgtgcgtggaaagcgtagacaaggagatgcaggtattggtgagtc<br>ggatcgcagcttggatggccacttacctgaatgaccacctagagccttggattcagg<br>agaacggcggctgggatacttttgtggaactctatgggaacaatgcagcagccgag<br>agccgaaagggccaggaacgcggatccggtggcggcagcggcggagggtccg<br>gaggcgacagcttctggatcttcgtgaagagactgtgaatggccacaaccatgcgat<br>cgtacccatacgatgttccagattacgctgaattcatggcgcgtaaggtcgatctcacc<br>tcctgcgatcgcgagccgatccacatccccggcagcattcagccgtgcggctgccta<br>ctagcctgcgacgcgcaggcggtgcggatcacgcgcattacggaaaatgccggcg<br>cgttctttggacgcgaaactccgcgggtcggtgagctactcgccgattacttcggcga<br>gaccgaagcccatgcgctgcgcaacgcactggcgcagtcctccgatccaaagcg<br>accggcgctgatcttcggttggcgcgacggcctgaccggccgcaccttcgacatctc<br>actgcatcgccatgacggtacatcgatcatcgagttcgagcctgcggcggccgaac<br>aggccgacaatccgctgcggctgacgcggcagatcatcgcgcgcaccaaagaac<br>tgaagtcgctcgaagagatggccgcacgggtgccgcgctatctgcaggcgatgctc<br>ggctatcaccgcgtgatgttgtaccgcttcgcggacgacggctccgggatggtgatc<br>ggcgaggcgaagcgcagcgacctcgagagctttctcggtcagcactttccggcgtc<br>gctggtcccgcagcaggcgcggctactgtacttgaagaacgcgatccgcgtggtctc<br>ggattcgcgcggcatcagcagccggatcgtgcccgagcacgacgcctccggcgcc<br>gcgctcgatctgtcgttcgcgcacctgcgcagcatctcgccctgccatctcgaatttctg<br>cggaacatgggcgtcagcgcctcgatgtcgctgtcgatcatcattgacggcacgctat<br>ggggattgatcatctgtcatcattacgagccgcgtgccgtgccgatggcgcagcgcg<br>tcgcggccgaaatgttcgccgacttcttatcgctgcacttcaccgccgcccaccacca<br>acgctaa |
| pAG1976 | CFAST2-LaminA/C-IRES- HA-iRFP670 | atgggcgacagcttctggatcttcgtgaagagactgtccggaggaggcggcagcgg<br>cggagggggatccgagaccccgtcccagcggcgcgccacccgcagcggggcgc<br>aggccagctccactccgctgtcgcccacccgcatcacccggctgcaggagaagga<br>ggacctgcaggagctcaatgatcgcttggcggtctacatcgaccgtgtgcgctcgctg<br>gaaacggagaacgcagggctgcgccttcgcatcaccgagtctgaagaggtggtca<br>gccgcgaggtgtccggcatcaaggccgcctacgaggccgagctcggggatgccc<br>gcaagacccttgactcagtagccaaggagcgcgcccgcctgcagctggagctgag<br>caaagtgcgtgaggagtttaaggagctgaaagcgcgcaataccaagaaggaggg<br>tgacctgatagctgctcaggctcggctgaaggacctggaggctctgctgaactccaa<br>ggaggccgcactgagcactgctctcagtgagaagcgcacgctggagggcgagct<br>gcatgatctgcggggccaggtggccaagcttgaggcagccctaggtgaggccaag<br>aagcaacttcaggatgagatgctgcggcgggtggatgctgagaacaggctgcaga<br>ccatgaaggaggaactggacttccagaagaacatctacagtgaggagctgcgtga<br>gaccaagcgccgtcatgagacccgactggtggagattgacaatgggaagcagcgt<br>gagtttgagagccggctggcggatgcgctgcaggaactgcgggcccagcatgagg<br>accaggtggagcagtataagaaggagctggagaagacttattctgccaagctgga<br>caatgccaggcagtctgctgagaggaacagcaacctggtgggggctgcccacga<br>ggagctgcagcagtcgcgcatccgcatcgacagcctctctgcccagctcagccagc<br>tccagaagcagctggcagccaaggaggcgaagcttcgagacctggaggactcac<br>tggcccgtgagcgggacaccagccggcggctgctggcggaaaaggagcgggag<br>atggccgagatgcgggcaaggatgcagcagcagctggacgagtaccaggagctt |

| | | |
|---|---|---|
| | | ctggacatcaagctggccctggacatggagatccacgcctaccgcaagctcttgga<br>gggcgaggaggagaggctacgcctgtcccccagccctacctcgcagcgcagccg<br>tggccgtgcttcctctcactcatcccagacacagggtgggggcagcgtcaccaaaa<br>agcgcaaactggagtccactgagagccgcagcagcttctcacagcacgcacgca<br>ctagcgggcgcgtggccgtggaggaggtggatgaggagggcaagtttgtccggct<br>gcgcaacaagtccaatgaggaccagtccatgggcaattggcagatcaagcgcca<br>gaatggagatgatcccttgctgacttaccgcttcccaccaaagttcaccctgaaggct<br>gggcaggtggtgacgatctgggctgcaggagctggggccacccacagcccccta<br>ccgacctggtgtggaaggcacagaacacctggggctgcgggaacagcctgcgta<br>cggctctcatcaactccactggggaagaagtggccatgcgcaagctggtgcgctca<br>gtgactgtggttgaggacgacgaggatgaggatggagatgacctgctccatcacca<br>ccacggctcccactgcagcagctcgggggaccccgctgagtacaacctgcgctcg<br>cgcaccgtgctgtgcgggacctgcgggcagcctgccgacaaggcatctgccagcg<br>gctcaggagcccaggtgggcggacccatctcctctggctcttctgcctccagtgtcac<br>ggtcactcgcagctaccgcagtgtgggggcagtgggggtggcagcttcggggac<br>aatctggtcacccgctcctacctcctgggcaactccagcccccgaacccagagccc<br>ccagaactgcagcatcatgtaaatggccacaaccatgcgatcgtacccatacgatgt<br>tccagattacgctgaattcatggcgcgtaaggtcgatctcacctcctgcgatcgcgag<br>ccgatccacatccccggcagcattcagccgtgcggctgcctactagcctgcgacgc<br>gcaggcggtgcggatcacgcgcattacggaaaatgccggcgcgttctttggacgcg<br>aaactccgcgggtcggtgagctactcgccgattacttcggcgagaccgaagcccat<br>gcgctgcgcaacgcactggcgcagtcctccgatccaaagcgaccggcgctgatctt<br>cggttggcgcgacggcctgaccggccgcaccttcgacatctcactgcatcgccatga<br>cggtacatcgatcatcgagttcgagcctgcggcggccgaacaggccgacaatccg<br>ctgcggctgacgcggcagatcatcgcgcgcaccaaagaactgaagtcgctcgaag<br>agatggccgcacgggtgccgcgctatctgcaggcgatgctcggctatcaccgcgtg<br>atgttgtaccgcttcgcggacgacggctccgggatggtgatcggcgaggcgaagcg<br>cagcgacctcgagagctttctcggtcagcactttccggcgtcgctggtcccgcagca<br>ggcgcggctactgtacttgaagaacgcgatccgcgtggtctcggattcgcgcggcat<br>cagcagccggatcgtgcccgagcacgacgcctccggcgccgcgctcgatctgtcgt<br>tcgcgcacctgcgcagcatctcgccctgccatctcgaatttctgcggaacatgggcgt<br>cagcgcctcgatgtcgctgtcgatcatcattgacggcacgctatggggattgatcatct<br>gtcatcattacgagccgcgtgccgtgccgatggcgcagcgcgtcgcggccgaaatg<br>ttcgccgacttcttatcgctgcacttcaccgccgcccaccaccaacgctaa |
